# A Weight Loss Fibrotic Response in Visceral Fat Is Driven by Defective Lysosome Dependent Collagenolysis

**DOI:** 10.64898/2026.09.29.755405

**Authors:** Clémentine Rebière, Amanda Oliveira, Sébastien Dussaud, Iman Haddad, Agatha de Assis-Ferreira, Adma Babêtto, Christine Rouault, Fatiha Merabtene, Cheima Benbida, Jimon Boniface Abatan, Simone Vargas da Silva, Lais Bhering Martins, Licia Torres, Ana Letícia Malheiros Silveira, Cindy Rose, Laurent Genser, Fabrizio Andreelli, Isabelle Dugail, Marina Chaves de Oliveira, Joelle Vinh, Simon Lecoutre, Adaliene Versiani Matos Ferreira, Karine Clément, Emmanuel L. Gautier, Genevieve Marcelin

**Affiliations:** Sorbonne Université, INSERM, Nutrition and obesities: systemic approach research group, Nutriomics; Paris F-75013, France; Immunometabolism, Department of Nutrition, Nursing School, Universidade Federal de Minas Gerais; Belo Horizonte 31270-901, Brazil; ESPCI Paris, PSL University, UAR CNRS 2051, Biological Mass Spectrometry and Proteomics; Paris, France; Cellular and Molecular Pharmacology, Department of Cell Biology, Institute of Biology, Universidade do Estado do Rio de Janeiro; Rio de Janeiro 20551-030, Brazil; Assistance Publique Hôpitaux de Paris, Visceral Surgery Department, Pitié-Salpêtrière Hospital; Paris, France; Assistance Publique Hôpitaux de Paris, Diabetology Department, Pitié-Salpêtrière Hospital; Paris, France; Assistance Publique Hôpitaux de Paris, Nutrition Department, CRNH Ile de France, Pitié-Salpêtrière Hospital; Paris, France

## Abstract

Obesity induces pathological remodeling of white adipose tissue (WAT), including chronic inflammation and fibrosis, which drive metabolic dysfunction. Although weight loss (WL) is the cornerstone of obesity treatment and improves systemic metabolic health, its impact on WAT remodeling remains incompletely understood. Using a mouse model of diet-induced obesity followed by nutritional weight loss, we show that visceral WAT (vWAT) fails to fully recover despite normalization of body weight and marked metabolic improvment. Notably, vWAT fibrosis persists and continues to progress after WL.

This occurs while extracellular matrix (ECM) production by adipose progenitor cells is markedly reduced following WL, explaining why pharmacological inhibition of ECM synthesis fails to prevent fibrosis during WL. Instead, we observed impaired collagen degradation by both progenitor cells and macrophages, along with lysosomal dysfunction, during WL. This suggests that defective ECM degradation, rather than continued ECM production, underlies fibrosis persistence and aggravation during WL. Together, these findings identify impaired collagenolysis as a critical barrier to adipose tissue recovery after WL and suggest that enhancing ECM clearance, rather than inhibiting ECM synthesis, may represent a more effective therapeutic strategy to restore adipose tissue function.

## Introduction

Loss of white adipose tissue (WAT) homeostasis is a central driver of obesity-associated metabolic disorders, including insulin resistance, hepatic steatosis and type 2 diabetes (1, 2). Among the key alterations contributing to WAT dysfunction is fibrosis, characterized by excessive accumulation of extracellular matrix (ECM) components that restrict WAT plasticity and impair its metabolic and endocrine functions (3, 4). During obesity progression, ECM accumulation is tightly linked to chronic inflammatory activation (4, 5) and the expansion of pro-fibrotic stromal populations, including fibro-inflammatory progenitors (FIPs) (2, 6, 7). Consistent with this, inhibition of these pathways mitigates WAT fibrosis during obesity (7–9). In parallel, macrophages contribute to ECM remodeling by regulating inflammatory responses and collagen turnover (10–13). In humans, WAT fibrosis correlates with insulin resistance and an increased risk of type 2 diabetes, underscoring the importance of maintaining ECM homeostasis for metabolic health (2, 4, 9, 14–18).

Weight loss (WL), achieved through lifestyle changes, pharmacotherapy or bariatric surgery remains the primary therapeutic strategy for obesity management. Sustained reduction in body weight (10% or more) is associated with significant improvements in metabolic health (19, 20). Visceral fat, whose abundance and functional activity are strongly associated with cardiometabolic risk, is significantly reduced by these interventions (21). However, accumulating clinical and experimental evidence indicates that structural remodeling of the tissue is incompletely reversed (22, 23). Persistent fibrosis has thus emerged as a hallmark of post-weight-loss WAT, yet the mechanisms responsible for its maintenance remain poorly understood (22, 24).

Here, we show that the initial phase of WL (before weight stabilization) fails to restore ECM homeostasis despite substantial metabolic improvement. We demonstrate that visceral white adipose tissue (vWAT) exhibits persistent and even progressive fibrosis during WL, despite resolution of obesity-associated inflammatory and pro-fibrotic transcriptional programs. Mechanistically, fibrosis persistence is associated with defective collagen degradation by both adipose progenitor cells and macrophages, accompanied by lysosomal dysfunction. Accordingly, pharmacological inhibition of ECM synthesis fails to prevent fibrosis after WL.

Together, our findings support a model in which ECM deposited during obesity is insufficiently resolved following weight reduction due to impaired cellular collagenolysis rather than continued ECM production. These results suggest that targeting collagen turnover, rather than ECM production, during the metabolically favorable window of WL may represent a more effective strategy to restore WAT function.

## Results

### Visceral WAT dysfunctions persist after weight loss post-obesity

Weight loss (WL) remains the cornerstone of obesity management. However, accumulating evidence suggests that vWAT does not fully recover its functional integrity after obesity. In particular, although sustained impairments in post-obese vWAT resilience have been described (22–24), the mechanisms underlying the persistence of vWAT fibrosis following WL remain largely unexplored. To address this question, we used the C3H mouse strain, a well-established model of obesity-associated fibrosis characterized by robust extracellular matrix deposition in the vWAT (8, 9). This model provides a unique opportunity to investigate the mechanisms governing fibrosis persistence and extracellular matrix remodeling following weight loss. We therefore performed an in-depth characterization of vWAT remodeling and function during the post-obese state.

To this end, we analyzed C3H mice fed a HFD for 4 weeks, followed by 4 weeks on a chow diet to induce WL (Formerly obese, FOb) (**Figure 1A**). FOb animals were compared to age-matched lean (Le) animals and to baseline obese mice before WL initiation (Ob 4w ; **Figure 1A**). Following the nutritional switch, body weight, body composition and fat depot masses in FOb mice normalized (**Figure 1B-E**, **S1A**). Accordingly, adipocyte size in both depots returned to levels observed in lean animals (**Figure 1F**, **S1B–C**) and glucose tolerance improved, reaching values comparable to those of lean mice (**Figure 1G**), contrasting with the impaired glucose clearance seen in Ob 4w animals (**Figure S1D, E**).

**Figure 1.**
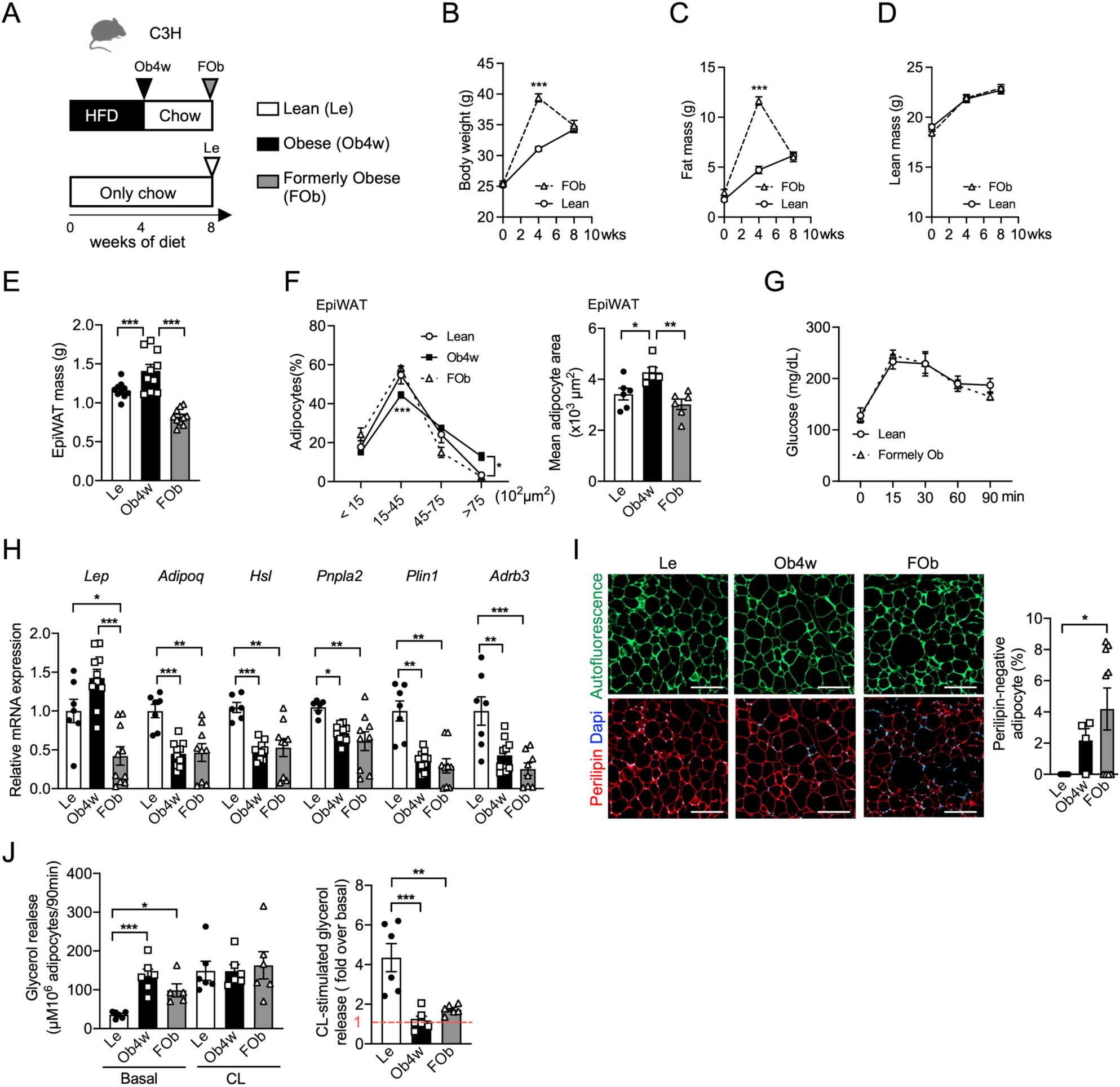
EpiWAT exhibited alterations in formerly obese mice. (**A**) Experimental design showing dietary interventions (chow or high-fat diet, as indicated) and the timeline for obesity induction and weight loss. Arrows indicate the feeding durations for lean (Le), obese (Ob4w) and formerly obese (FOb) groups. (**B**) Body weight, (**C**) fat mass, (**D**) lean mass curves and (**E**) EpiWAT mass in Le, Ob4w and FOb mice (n = 10). (**F**) Adipocyte size distribution and mean adipocyte area in EpiWAT of Le, Ob4w and FOb mice (n = 6). (**G**) Glucose tolerance test (GTT) in age-matched Le and FOb mice after 8 weeks of dietary intervention (n = 6). (**H**) Relative mRNA expression of adipocyte markers in EpiWAT measured by quantitative PCR (n = 7–10). (**I**) Representative immunofluorescence staining of EpiWAT sections from Le, Ob4w and FOb mice with anti-Perilipin 1 (red) and DAPI (blue); tissue autofluorescence (green) was used to assess structural integrity (scale bar, 200 μm). The percentage of Perilipin-negative adipocytes was quantified in EpiWAT sections (n = 4–9). (**J**) Left, basal and CL-316,243 (CL)-stimulated glycerol release from EpiWAT explants, normalized to adipocyte number in Le, Ob4w and FOb mice (n = 6). Right, fold change in CL-induced lipolysis over basal glycerol release in EpiWAT explants from Le, Ob4w and FOb mice (n = 6). (**B-D**, **G**) Statistical significance was determined by Student’s t-test ; (**E, F, H–J**) one-way ANOVA followed by Tukey’s multiple comparisons test.

In addition to this normalization of fat mass and glucose homeostasis following WL, we investigated whether vWAT function was also restored. We first assessed molecular and cellular alterations in vWAT (epidydimal fat depot, EpiWAT) following WL. Gene expression analysis revealed, except for *Lep*, persistent downregulation of key adipocyte genes (*Adipoq*, *Hsl, Pnpla2*, *Plin1* and *Adrb3*) in EpiWAT of FOb mice compared to lean mice (**Figure 1H**). These defects were specific to EpiWAT, as the subcutaneous WAT (inguinal fat depot, IngWAT) showed no significant changes for these genes in FOb compared to lean mice (**Figure S1F**). Immunofluorescent staining further revealed a reduction in Perilipin-1 expression in adipocytes from FOb animals compared with lean mice, which was even more pronounced than in Ob4w mice (**Figure 1I**). Together with the reduced *Plin1* transcript levels (**Figure 1H**), these findings are consistent with persistent adipocyte dysfunction following WL. To determine whether these defects translated into functional abnormalities, we next examined the lipolytic capacity of adipocytes. EpiWAT from FOb mice exhibited persistently elevated basal lipolysis (**Figure 1J**), a hallmark feature of adipocyte dysfunction in obesity (25). Moreover, β3-adrenergic receptor–stimulated lipolysis, critical for catecholamine-driven lipid mobilization in mice (26), remained blunted (**Figure 1J**), consistent with sustained downregulation of *Adrb3*, which encodes β3-AR (**Figure 1H**). Again, these defects were specific to EpiWAT, as IngWAT showed no significant alterations in stimulated lipolysis or *Adrb3* expression (**Figure S1F–H**).

In summary, despite systemic metabolic improvements, visceral EpiWAT in formerly obese mice retains significant molecular and functional defects after WL, highlighting a persistent impairment in adipose tissue recovery.

### Weight loss exacerbates fibrosis in visceral adipose tissue

As fibrosis contributes to WAT dysfunction (4, 27), we therefore examined the fibrotic remodeling after WL. Histological analysis revealed increased collagen deposition in the EpiWAT of FOb mice compared to Ob4w animals, with fibrosis worsening over the course of the WL period (**Figure 2A**). Hydroxyproline content, a biochemical proxy for total collagen (28), was quantified and normalized to either EpiWAT mass (**Figure 2B**) or measured in the entire fat pad (**Figure 2C**). Both approaches confirmed the increased fibrosis accumulation during WL. In contrast, no such changes were observed in IngWAT, indicating a depot-specific response (**Figure S1B**).

**Figure 2.**
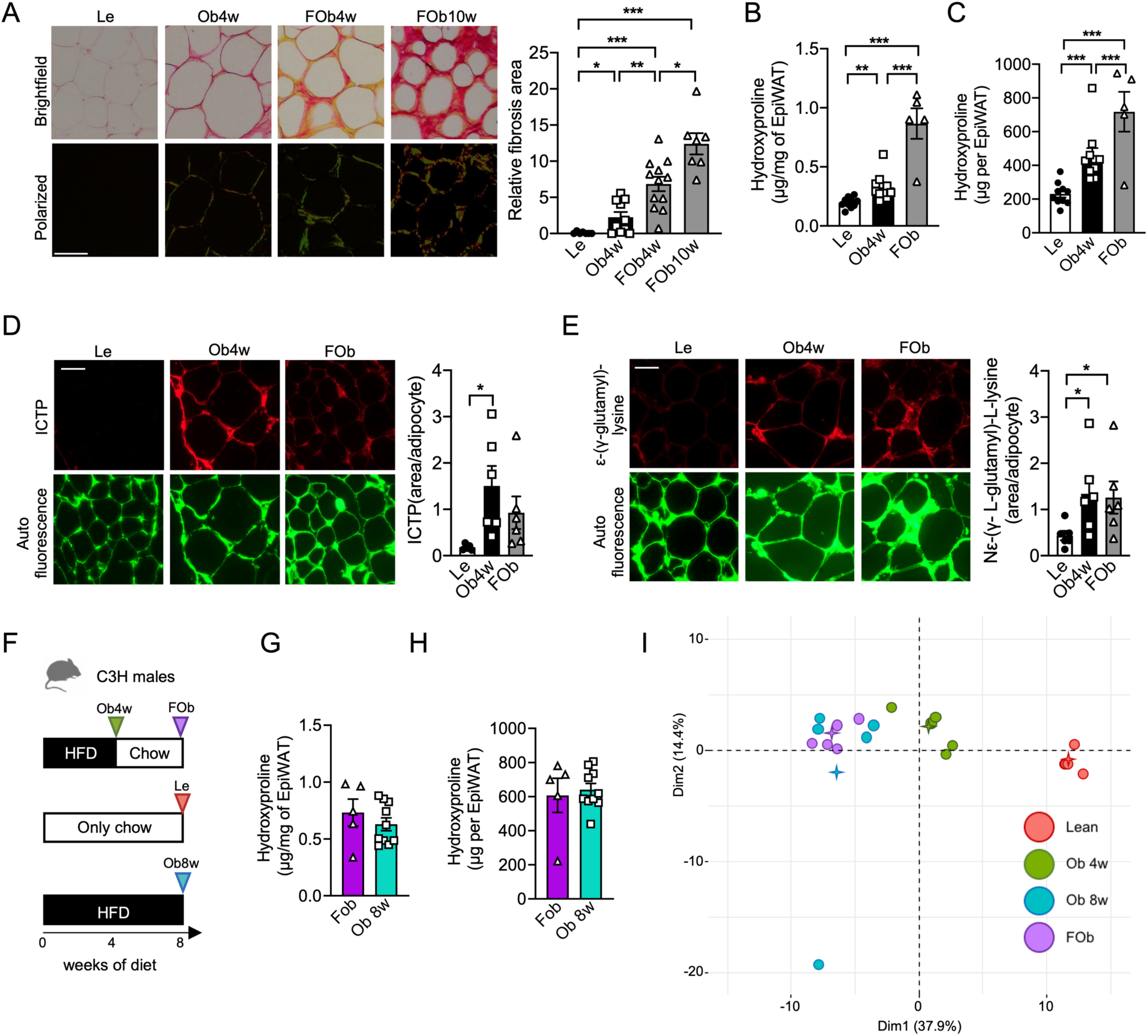
Fibrosis deposition in EpiWAT of FOb mice. (**A**) Representative picrosirius red-stained histological sections of EpiWAT from lean (Le), obese (Ob4w) and formerly obese (FOb) mice, imaged under brightfield and polarized light (scale bar, 100 μm). We analyzed two FOb groups, both fed a high-fat diet (HFD) for 4 weeks, then switched to a chow diet for either 4 weeks (named FOb or FOb 4w) or 10 weeks (FOb 10w). Quantification of fibrosis area normalized to adipocyte number (relative values; Lean group set to 1), based on picrosirius red staining (n=7-12). (**B**) Hydroxyproline content in EpiWAT normalized to fat pad mass (μg/mg) and (**C**) total hydroxyproline content per fat pad (n=5-10). (**D**) Representative immunofluorescence staining of EpiWAT sections from Le, Ob4w and FOb mice using an anti-ICTP antibody (red). Tissue autofluorescence (green) highlights structural integrity (scale bar, 50 μm). Quantification (arbitrary units, AU) is normalized to adipocyte number (n=5–6). (**E**) Representative immunofluorescence staining of EpiWAT using an anti-Nε-(γ-L-glutamyl)-L-lysine antibody (red); autofluorescence (green) highlights tissue integrity (scale bar, 50 μm). Quantification (AU) is normalized to adipocyte number (n=6). (**F**) Experimental design showing dietary interventions (chow or high-fat diet, as indicated) and the timeline for obesity induction and weight loss. Arrows indicate feeding durations for the groups: lean (Le, orange), formerly obese (FOb, purple) and obese mice fed HFD for 4 weeks (Ob4w, green) or 8 weeks (Ob8w, blue). (**G**–**H**) Fibrosis quantification in EpiWAT of FOb and Ob8w mice by measuring: (**G**) Hydroxyproline content normalized to fat pad mass (μg/mg) and (**H**) Total hydroxyproline content per fat pad (n=5–10). (**I**) Principal component analysis (PCA) of extracellular matrix (ECM) proteins from Le, Ob4w, Ob8w and FOb EpiWAT (n=5). (**A**–**E**) Statistical testing was performed using one-way ANOVA with Tukey’s multiple comparisons test. (**G**– **H**) Statistical significance was assessed using Student’s t-test.

While sexual dimorphism is well recognized in obesity, most studies have focused on differences in susceptibility to weight gain and metabolic dysfunction (29), whereas less attention has been paid to WAT fibrosis. In this setting, we analyzed female C3H mice subjected to the same experimental design than males (**Figure 1A**). Body weight, body composition and fat pad mass increased in HFD-fed females and were normalized following weight loss (**Figure S2A-E**), as observed in males. Female C3H mice displayed body weight and fat mass values comparable to males after 4 weeks of HFD feeding (**Figure S2A, B**). In females, 4 weeks of HFD feeding induced glucose intolerance (**Figure S2F**), which was normalized after 4 weeks of weight loss (**Figure S2G**). In agreement with our previous observations (9), obesity also increased hydroxyproline content in the vWAT of females (periovarian fat depot, ovaWAT), indicating the development of fibrosis (**Figure S2H, I**). However, the magnitude of fibrosis was lower than that observed in males (**Figure S2J**). Importantly, in contrast to our observations in male mice, we did not detect further fibrosis accumulation following weight loss in females. The apparent increased hydroxyproline content (**Figure S2H**), which was normalized to fat mass after weight loss, most likely reflected the reduction in ovaWAT mass (**Figure S2E**) rather than additional collagen deposition. Indeed, total hydroxyproline content per fat pad was similar between obese and post-obese females (**Figure S2I**). These findings suggest that weight loss does not promote additional fibrosis accumulation in ovaWAT.

Given that post-translational crosslinking of ECM proteins, such as collagens, critically shapes ECM architecture and contributes to fibrotic stiffness and tissue dysfunction (30), we next assessed two key crosslinking products, the carboxy-terminal cross-linked telopeptide of type I collagen (ICTP) and Nε-(γ-L-glutamyl)-L-lysine, by immunostaining (31, 32) in males. Both markers were significantly elevated in obese mice (Ob4w) compared to lean mice and remained high in FOb males (**Figure 2D**, **E**), indicating persistent ECM stiffening despite WL.

Because WAT fibrosis was minimal after only 4 weeks of HFD in males (**Figure 2A-C**), the subsequent collagen accumulation observed in FOb mice likely reflects active remodeling rather than residual fibrosis from early obesity. To further delineate the dynamics of ECM remodeling, we compared EpiWAT from lean males (8-week chow), Ob4w and Ob8w animals (4 and 8 weeks of HFD, respectively) and FOb mice (4-week HFD followed by 4-week chow, FOb) (**Figure 2F**). Hydroxyproline content in FOb and Ob8w mice was comparable (**Figure 2G, H**), suggesting that WL induces fibrotic changes of similar magnitude to those observed in advanced obesity.

To gain further insight into the ECM composition in these weight gain-weight loss conditions, we performed proteomic profiling of the EpiWAT matrisome. Mass spectrometry identified 158 matrisome proteins, including 68 core matrisome and 90 matrisome-associated proteins (33). Principal component analysis (PCA) revealed that samples from FOb and Ob8w mice clustered together, distinct from lean and Ob4w groups (**Figure 2I**), consistent with a convergent fibrotic signature driven by either sustained obesity or WL. Accordingly, no significant differences in individual protein levels were detected between FOb and Ob8w mice (**Figure S3**). Of the 158 identified matrisome proteins, 104 were differentially expressed in both FOb and Ob8w relative to lean (**Figure S3A**). Notably, 70% of these changes were shared between FOb and Ob8w mice (**Figure S3A, C**), while only 5.8% and 22.1% were specific to Ob8w and FOb, respectively (**Figure S3A, B, D**). Downregulated proteins included basement membrane components (e.g., LAMB1, COL4A1, NID1) and protease inhibitors, suggesting compromised matrix integrity. In parallel, FOb and Ob8w mice showed increased expression of cysteine proteases (CTSS, CTSB) and collagen-stabilizing enzymes (LOXL1, P4HA1), consistent with active ECM remodeling and fiber crosslinking.

Together, these findings reveal that EpiWAT fibrosis is not only retained but intensified following WL. Importantly, this remodeling is characterized by impaired basement membrane architecture and increased ECM crosslinking, potentially contributing to major persistent tissue dysfunction in the formerly obese state.

### Macrophage persistence and reprogramming in the EpiWAT after weight loss post-obesity

As proinflammatory immune cell recruitment contributes to obesity-induced fibrosis (4, 5, 8), we next assessed whether obesity-associated macrophage accumulation was maintained after WL. Flow cytometry analysis of EpiWAT revealed that total macrophage numbers (**Figure S4A, B**) were elevated approximately fourfold in both Ob4w and FOb mice compared to lean controls (**Figure 3A**). However, due to the reduced EpiWAT mass in FOb mice, macrophage density was significantly higher than in Ob4w mice (**Figure 3B**), indicating a proportionally greater macrophage burden after WL. To further define macrophage phenotypes, we sorted EpiWAT macrophages from lean, Ob4w and FOb mice for RNA sequencing. Prior studies have identified a distinct subpopulation of monocyte-derived macrophages, termed lipid-associated macrophages (LAMs), characterized by high CD9 expression and a transcriptional program enriched in lipid metabolism, lysosomal function and inflammatory pathways (34). Gene set enrichment analysis (GSEA) revealed that LAM-associated transcripts were enriched not only in Ob4w versus lean mice (data not shown), but also in FOb versus Ob4w macrophages (**Figure 3C**). Flow cytometry confirmed an increased proportion of CD9⁺ LAMs in FOb mice compared to Ob4w (**Figure 3D–F**), suggesting that the obesity-associated macrophage program is maintained in adipose tissue even after WL.

**Figure 3.**
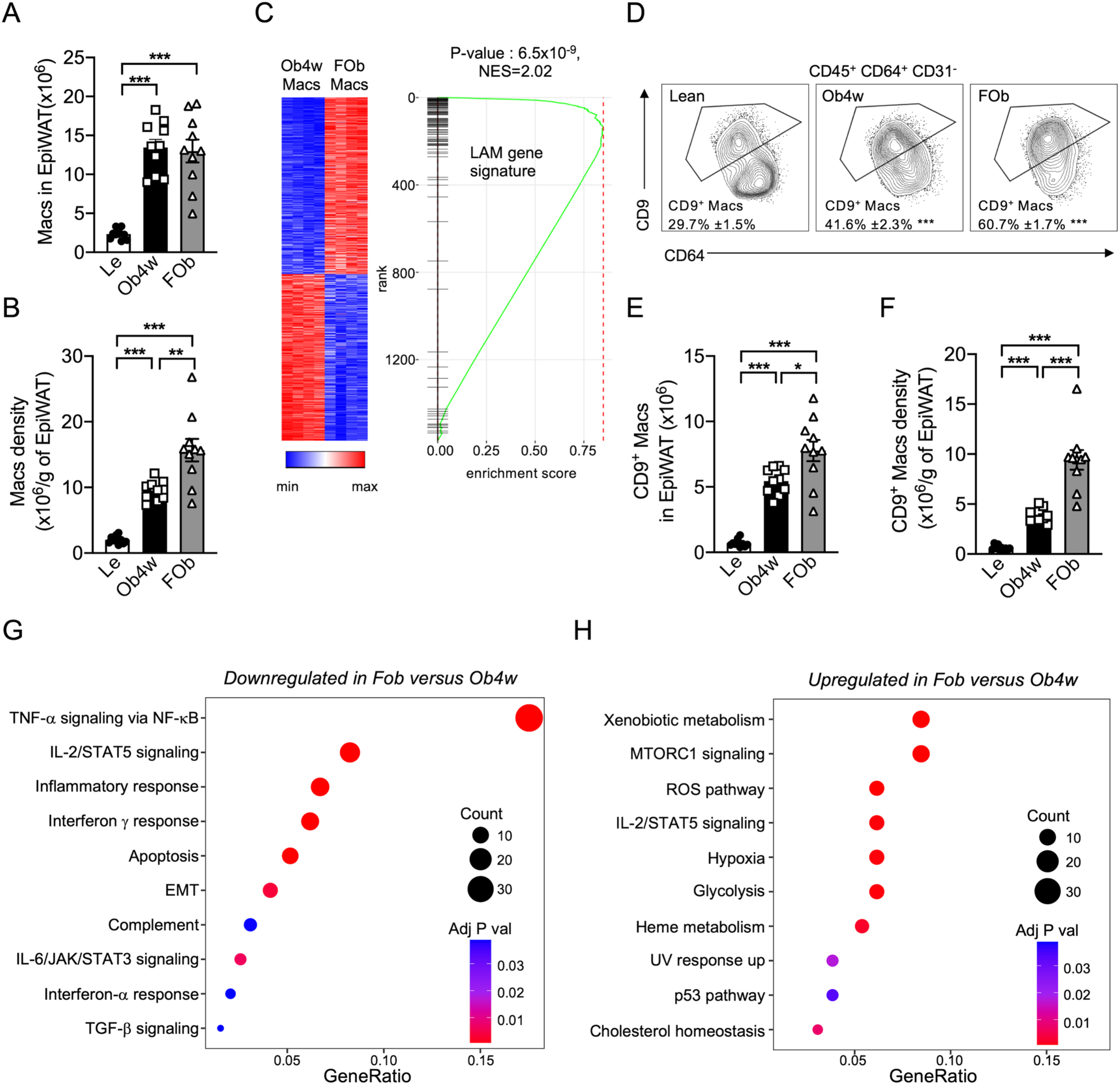
Lipid-Associated Macrophage Accumulation persisted in EpiWAT of FOb mice. (**A**) Total macrophage number (CD45⁺ CD64⁺ CD31⁻) in EpiWAT and (**B**) macrophage number per gram of EpiWAT from lean (Le), obese (Ob4w) and formerly obese (FOb) mice, quantified by flow cytometry (n = 10). (**C**) Gene set enrichment analysis (GSEA) of the lipid-associated macrophage (LAM) gene signature within the total macrophage population (CD45⁺ CD64⁺ CD31⁻) from Ob4w and FOb mice (n = 4). (**D**) Representative FACS plot and mean percentage (± SEM) of CD9⁺ macrophages (Macs) in EpiWAT from Le, Ob4w and FOb C3H mice (n=10). (**E**) Total CD9⁺ macrophage number in EpiWAT and (**F**) CD9⁺ macrophage number per gram of EpiWAT in Le, Ob4w and FOb mice, quantified by flow cytometry (n = 10). (**G**) Pathways enriched in macrophages isolated from Ob4w mice compared to those from FOb mice. (**H**) Pathways enriched in macrophages isolated from FOb mice compared to those from Ob4w mice. (**A**, **B**, **D**–**F**) Statistical significance was assessed using one-way ANOVA with Tukey’s multiple comparisons test. For panel D, statistical significance was defined as: a P<0.001 vs. Lean ; b P<0.001 vs Ob4w. (**C**) LIMMA analysis was used to identify differentially expressed genes, followed by GSEA of the CD9⁺ macrophage gene signature using Phantasus software. (**G**, **H**) Pathway enrichment was analyzed using ClusterProfiler, with false discovery rate (FDR) correction and adjusted p-values calculated using the Benjamini–Hochberg procedure.

TLR4 signaling in bone marrow-derived myeloid cells has previously been implicated in adipose tissue fibrosis in the C3H mouse strain (8). We therefore tested whether pharmacological inhibition of TLR4 might protect adipose tissue from fibrosis during WL. We administered TAK242—a TLR4 inhibitor with anti-inflammatory properties previously shown to reduce adipose tissue fibrosis in obese C3H mice (8), at the time of the diet switch (**Figure S5A**). TAK242 treatment had no effect on body weight or composition (**Figure S5B**), EpiWAT mass (**Figure S5C**) or collagen content (**Figure S5D–E**). Thus, fibrosis persistence after WL occurs independently from TLR4 signaling.

We next examined the transcriptional landscape of adipose tissue macrophages after WL using RNA sequencing. This revealed a marked shift in macrophage gene expression in FOb animals. Compared with Ob4w, FOb macrophages showed reduced inflammatory activation and antigen presentation, as indicated by downregulation of genes in the TNFA/NF-KappaB signaling pathway (**Figure 3G**; **Figure S6**), as well as genes encoding MHC class II components, including *H2-Ab1* (FC=0.33, P=5.10^-7^), *H2-Eb1* (FC=0.30, P=2.10^-6^) and *Cd74* (FC=0.34, P=1.10^-6^). In addition, FOb macrophages displayed increased expression of genes associated with oxidative and metabolic stress (*Hmox1*, *Gclc*, *Gclm*, *Txn1*, *Txnrd1*, *Cat*, *Srxn1*) and glycolysis and one-carbon metabolism (*Slc2a1*, *Hk2*, *Aldoa*, *Pkm*, *Mthfd2*) compared to Ob4w (**Figure 3H**). Finally, several immunoregulatory molecules (*Cd5l*, *Havcr2*, *Lilrb4a*, *Adgre5*) were enriched in FOb macrophages (**Figure S6).**

Together, these data indicate that adipose tissue macrophages undergo transcriptional reprogramming after WL, which is characterized by attenuation of pro-inflammatory and antigen-presenting programs, and the acquisition of stress-adapted and immunomodulatory features. This shift in FOb macrophage transcriptional state further suggests that specific mechanisms may underlie weight loss–associated adverse adipose tissue remodeling.

### Weight Loss Uncouples ECM production from fibrosis in Visceral Fat

We next evaluated a dual anti-fibrotic strategy targeting tyrosine kinase and COX2 signaling, using nintedanib (35, 36) and celecoxib (37), agents with established efficacy in EpiWAT fibrosis in our previous study (7). Co-administration of these compounds similarly failed to reduce collagen accumulation during WL (**Figure S5F–J**), mirroring the inefficacy of TAK242. To elucidate the mechanisms underlying the differential sensitivity to antifibrotic therapies between obesity-associated and WL–associated fibrosis, we analyzed ECM-related gene expression in EpiWAT. We aimed to uncover molecular differences in fibrotic remodeling that may influence treatment efficacy in these two physiologically distinct contexts. While these ECM-related genes were upregulated in Ob4w mice, they were downregulated in FOb animals, reaching levels comparable to those in lean mice (**Figure 4A-B**). Thus, despite the persistent and even increased fibrosis observed in the FOb EpiWAT (**Figure 2A-C**), the transcriptional program driving ECM synthesis was suppressed after WL. This likely explains the failure of TAK242 and the nintedanib-celecoxib combination in this context as WL itself is sufficient to suppress ECM gene expression, making further inhibition innefective.

**Figure 4.**
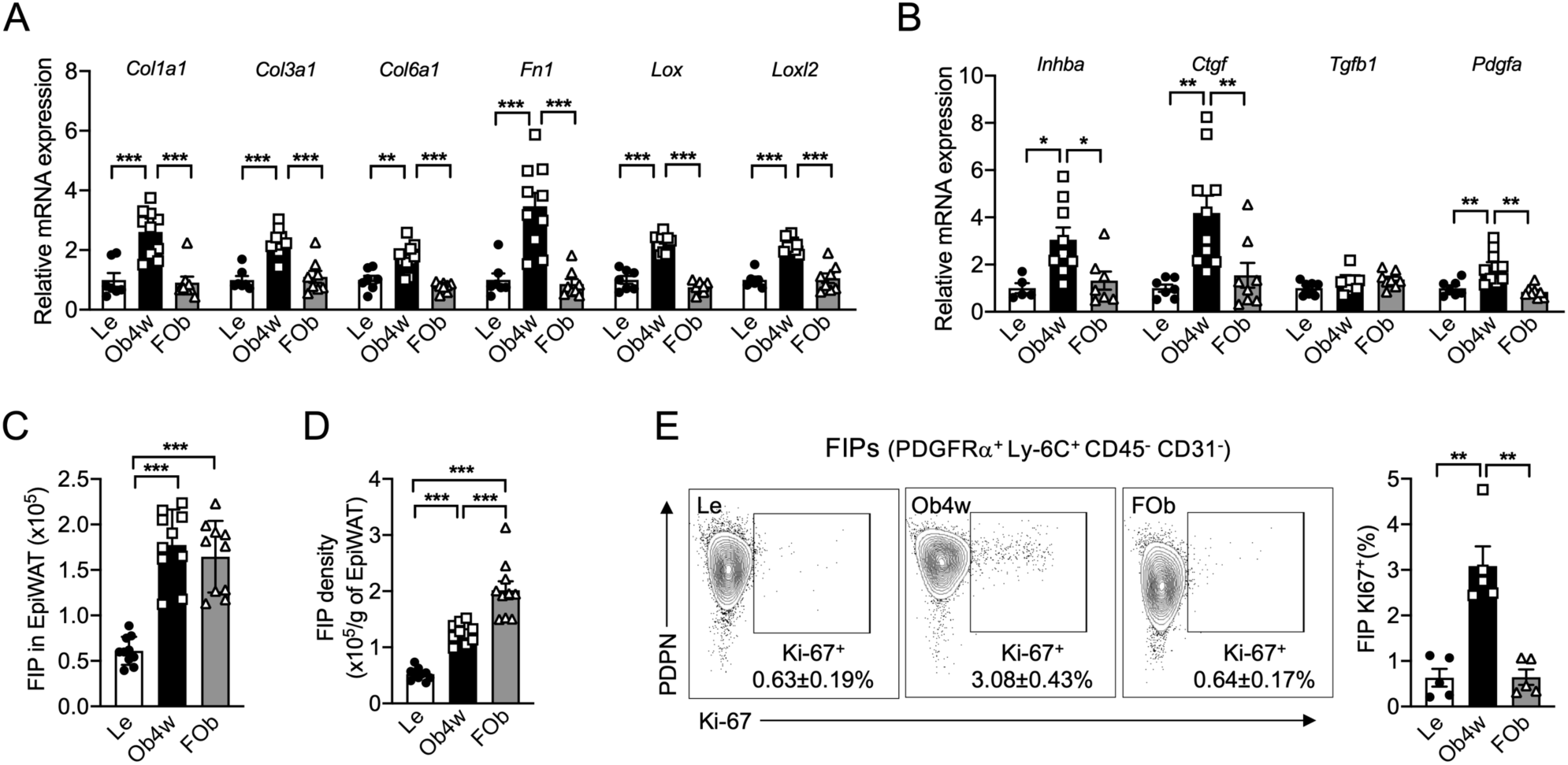
Dissociation of the tissue fibrosis signature from fibro-inflammatory progenitor (FIP) persistence. (**A**, **B**) Relative mRNA expression of (**A**) fibrosis-related ECM components and (**B**) fibrogenic cytokines measured by quantitative PCR in EpiWAT of lean (Le), obese (Ob4w) and formerly obese (FOb) C3H mice (n=5–10). (**C**) Total FIP (PDGFRα⁺ PDPN⁺ CD9⁺ Ly-6C⁺ CD45⁻ CD31⁻) count in EpiWAT (n=10) and (**D**) FIP count per gram of EpiWAT (n=10) quantified by flow cytometry. (**E**) Left: representative FACS plots showing Ki-67 expression in FIPs from EpiWAT stromal vascular fraction (SVF) of Le, Ob4w and FOb C3H mice. Right: percentage (mean+/-SEM) of Ki-67⁺ FIPs in EpiWAT (n=5). (**A**–**E**) Statistical testing was performed using one-way ANOVA with Tukey’s multiple comparisons test.

We next turned our attention to fibro-inflammatory progenitors (FIPs), a specific subset of CD9⁺ Ly-6C⁺ PDGFRα⁺ adipose stromal progenitor cells (**Figure S4A, C**) that expand during obesity and acquire pro-fibrotic and pro-inflammatory features (6, 9). Given their central role in obesity-associated fibrosis through increased extracellular matrix production (6, 9), we investigated whether the activation of this population was modulated after WL. Flow cytometry revealed elevated FIP abundance in both Ob4w and FOb mice relative to lean animals (**Figure 4C**), with FIP density (cell count per gram of fat) further increased in FOb animals due to adipose mass reduction (**Figure 4D**). FIP proliferation was enhanced in Ob4w mice but returned to baseline after WL (**Figure 4E**), probably indicating a halt in expansion although FIPs persisted and became more densely concentrated within EpiWAT of FOb mice. To further characterize FIP states in these different contexts, we performed transcriptomic profiling of sorted FIPs from lean, Ob4w and FOb mice. Principal component analysis revealed that FIPs from FOb mice clustered separately from both lean and Ob4w mice (**Figure 5A**). K-means clustering of differentially expressed genes between lean and FOb FIPs identified five distinct modules (**Figure 5B**). Cluster 1 included genes encoding ECM proteins, which were selectively downregulated in FOb FIPs compared to both lean and Ob4w animals. Cluster 2 comprised genes involved in epithelial–mesenchymal transition (EMT) and ECM remodeling, which were upregulated in Ob4w and FOb mice but attenuated in FOb relative to Ob4w, reflecting the tissue-level suppression of fibrotic markers (**Figure 4A**, **B**). These two clusters indicate that the fibrogenic activation of FIPs is significantly reduced following WL. To quantify this reversal, we focused on 46 EMT-related genes upregulated in Ob4w versus lean FIPs. Of these, 32 were significantly downregulated in FOb versus Ob4w and 15 returned to lean levels (**Figure 5C**), demonstrating broad suppression of the fibrogenic program within FIPs after WL. Cluster 3 included inflammatory genes, which were upregulated in both Ob4w and FOb FIPs compared to lean mice, suggesting persistent inflammatory activation. Cluster 4, linked to BMP signaling and ECM organization, was downregulated in Ob4w and remained suppressed in FOb mice. Notably, Cluster 5 enriched for NRF2 signaling, glutathione metabolism and lysosomal function, showed stronger induction in FOb than in Ob4w FIPs, pointing to elevated oxidative stress after WL.

**Figure 5.**
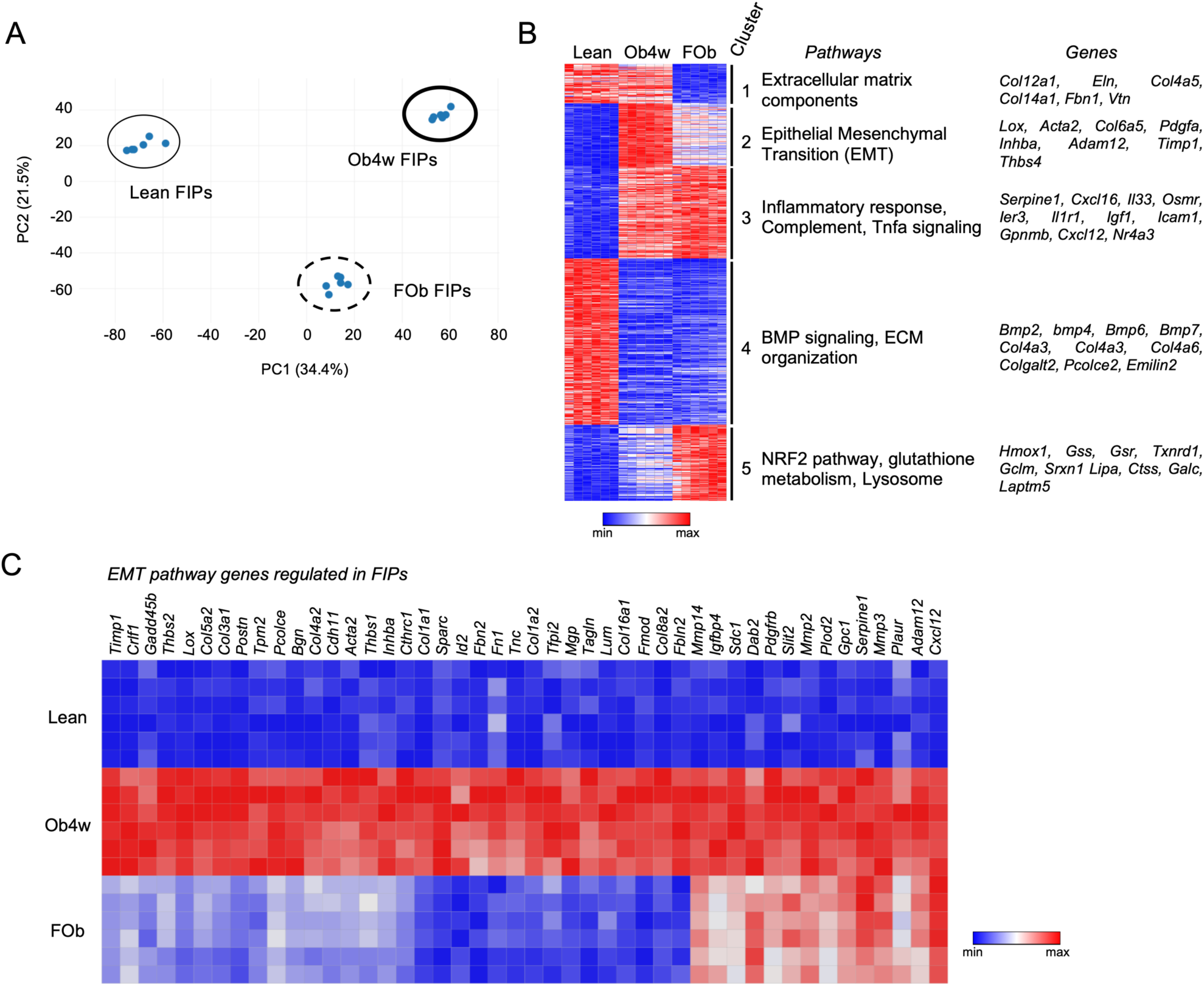
The fibro-inflammatory progenitors (FIPs) revert to an inactive phenotype during weight loss. (A) Principal component analysis (PCA) of FIP transcriptomes from Le, Ob4w and FOb mice (n=6). (B) K-means clustering analysis of FIPs from Le, Ob4w and FOb mice identified five clusters. Representative genes and enriched pathways for each cluster are shown (n=6). (**C**) Heatmap showing expression of genes in the epithelial-to-mesenchymal transition (EMT) pathway that are upregulated in FIPs from Ob4w vs. Le mice (fold change > 1.5, adjusted p-value < 0.05; n=6). (**B**, **C**) A Limma analysis was conducted to select differentially expressed genes between (**B**) Le and FOb and (**C**) Le and Ob4w (fold change > 1.5, adjusted p-value for multiple testing).

In summary, these findings reveal a striking uncoupling between ECM gene expression and ECM protein accumulation in the EpiWAT of formerly obese mice. Despite continued collagen deposition, the transcriptional program within FIPs that drives ECM synthesis is largely silenced after WL. This decoupling likely underlies the inefficacy of therapies targeting ECM production (7, 8) and suggests that post-obesity fibrosis is maintained by mechanisms beyond the production of ECM components. We therefore hypothesized that the incremental fibrosis observed during WL may arise from impaired matrix turnover or clearance, rather than ongoing ECM production.

### Lysosomal stress links impaired collagen degradation to fibrosis during weight loss

To investigate whether defective collagen degradation contributes to the worsening of EpiWAT fibrosis during WL, we developed an *ex vivo* assay using the DQ^TM^ collagen I, a fluorogenic substrate that emits FITC fluorescence upon proteolytic cleavage (38, 39). While FIPs are central to fibrosis development through enhanced ECM production, the entire progenitor cell population could participate to collagen degradation and fibrosis resolution after weight loss. We therefore did not restrict the analysis to the FIP subset, as ECM degradation and turnover are not necessarily confined to this population.

In lean EpiWAT, collagenolytic activity, as indicated by FITC fluorescence, was primarily observed in macrophages (CD45⁺ CD64⁺) and progenitors (PDGFRα⁺), while endothelial cells (CD31⁺ CD45⁻ PDGFRα⁻) showed minimal signal (**Figure 6A, B**). When comparing lean, Ob4w and FOb mice, both macrophages and progenitors from Ob4w and FOb mice exhibited significantly reduced collagen-degrading capacities relative to lean mice (**Figure 6C, D**), indicating that collagenolysis is impaired during obesity and remains defective after WL. We next assessed collagenolytic activity in macrophages and progenitor cells from IngWAT in males (**Figure S7 A, B**) and periovarian WAT (OvaWAT) in females (**Figure S7 C, D**). Unlike EpiWAT in males, neither depot exhibited alterations in collagen degradation capacity following obesity or weight loss, with comparable collagenolytic activity observed in Lean, Ob4w and FOb mice (**Figure S7 A-D**).

**Figure 6.**
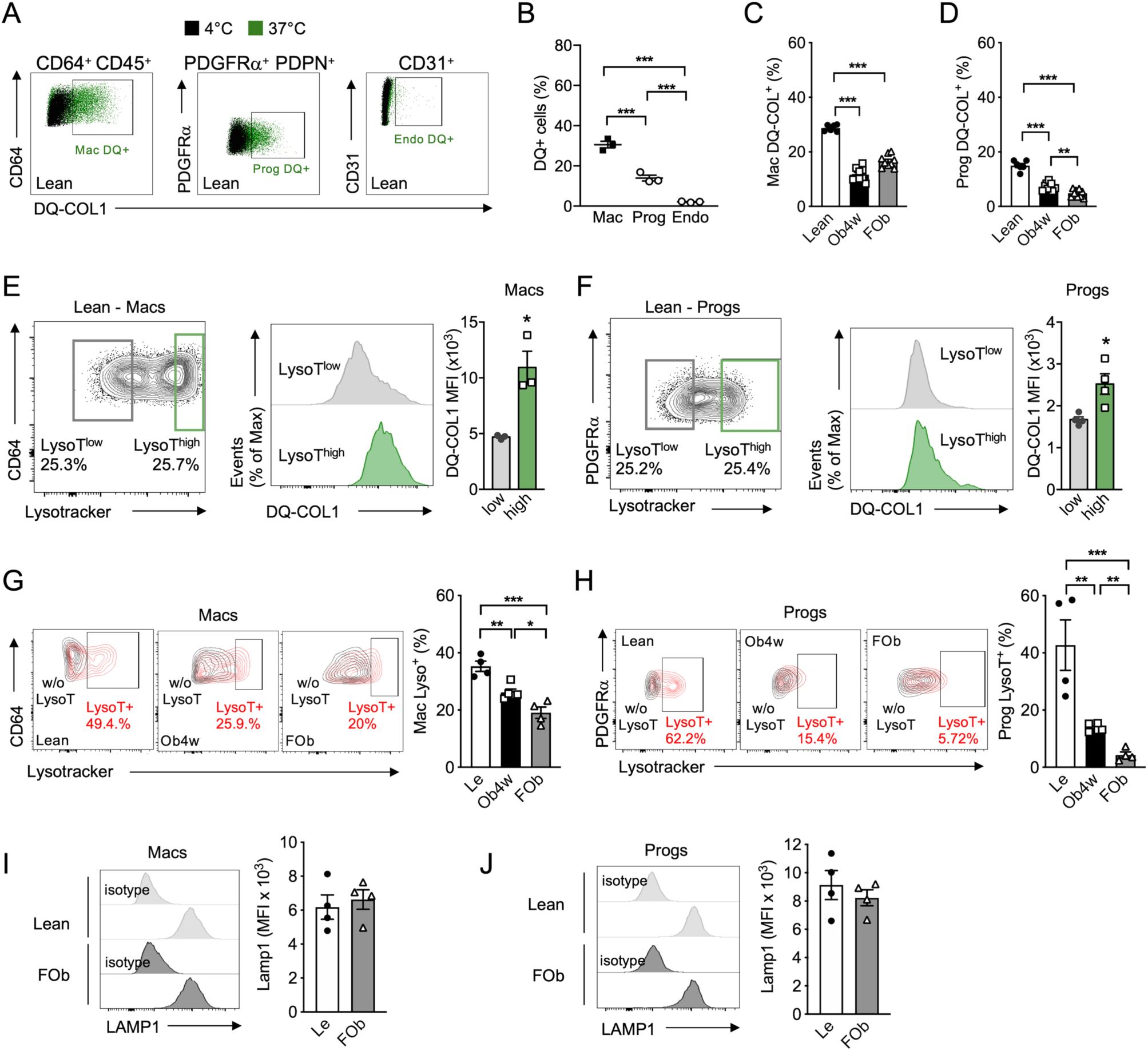
Reduced collagen 1 degradation and decreased lysosome signal in Macs and Progs of FOb C3H mice. (**A**) Representative FACS plots showing DQ-COL1⁺ macrophages (Macs), adipose progenitors (PDGFRα⁺; Progs) and endothelial cells (Endo) incubated at 4 °C (negative control, black) or 37 °C (green). (**B**) Mean percentage of DQ-COL1⁺ cells among Macs, Progs and endothelial cells (n=3). (**C**, **D**) Percentage of DQ-COL1⁺ cells in (**C**) Macs and (**D**) Progs isolated from EpiWAT of lean (Le), obese (Ob4w) and formerly obese (FOb) C3H mice (n=6–9). (**E**) Representative FACS plot showing the gating strategy to identify LysoTlow and LysoThigh Macs (based on Lysotracker staining) from lean EpiWAT and corresponding DQ-COL1 fluorescence intensity (n=3). (**F**) Same analysis as in (**E**) for Progs (n=3). (**G**) Quantification of LysoT⁺ Macs isolated from EpiWAT SVF of Le, Ob4w and FOb mice in the presence (red) or absence (black) of Lysotracker staining (n=4). (**H**) Quantification of LysoT⁺ Progs isolated from EpiWAT SVF of Le, Ob4w and FOb mice in the presence (red) or absence (black) of Lysotracker staining (n=4). (**I**) FACS analysis of LAMP1 expression in Macs from EpiWAT SVF of Le and FOb mice (n=4). (**J**) FACS analysis of LAMP1 expression in Progs from EpiWAT SVF of Le and FOb mice (n=4). (**B**–**D**, **G**, **H**) Statistical testing was performed using one-way ANOVA followed by Tukey’s multiple comparisons test. (**E**, **F**, **I**, **J**) Statistical significance was assessed using Student’s t-test.

To gain further insight into the alterations observed in EpiWAT, we analyzed the expression of genes encoding regulators of collagen degradation (40). At the whole-tissue level, the expression of these collagenolytic regulators remained largely unchanged across groups (**Figure S8A**). Yet, a few genes (e.g., *Mmp2*, *Mmp3*, *Mmp14*, *Timp1*) which were elevated during obesity, returned to baseline (i.e., lean) in FOb mice. Also, genes involved in collagen uptake were comparably expressed in FIPs (**Figure S8B**) and macrophages (**Figure S8C**) from lean and FOb mice, suggesting that uptake mechanisms are not responsible for the impaired collagenolysis.

We next examined whether lysosomal dysfunction could underlie this defect, as lysosomal pathways are essential for intracellular collagen degradation (39, 41). Revisiting the RNA sequencing data obtained from macrophages and FIPs isolated from the EpiWAT of lean, Ob4w and FOb mice, gene set enrichment analysis revealed significant enrichment of lysosome-associated genes (KEGG Lysosome pathway, 127 genes) in both macrophages (**Figure S9A-B**) and FIPs (**Figure S9C-D**) from Ob4w and FOb mice compared to lean animals. Given that lysosomal gene induction can reflect stress-induced compensatory responses (42), these findings point to potential lysosomal dysfunction.

To directly evaluate lysosomal function, we used LysoTracker, a pH-sensitive dye that accumulates in acidic lysosomes (43). Flow cytometry enables quantitative assessment of LysoTracker signal in large numbers of freshly isolated SVF cells, while minimizing manipulation and preserving their tissue-derived phenotype. In both macrophages and Progs, LysoTracker intensity strongly correlated with DQ™ collagen degradation (**Figure 6E-F**), confirming the requirement of lysosomal acidification for collagenolysis. Importantly, LysoTracker staining was reduced in Ob4w and further diminished in FOb mice in both macrophages (**Figure 6G**) and Progs (**Figure 6H**), indicating lysosomal alteration with obesity and WL. However, the lysosomal membrane protein LAMP1 (43), showed unchanged expression in macrophages and Progs from FOb animals compared to lean mice (**Figure 6I-J**), suggesting that lysosome abundance is preserved and that the defect lies specifically in acidification capacity. We also assessed LysoTracker staining in macrophages and progenitor cells from IngWAT in males (**Figure S7 E, F**) and ovaWAT in females (**Figure S7 G, H**). In agreement with the preserved collagen degradation capacity in these depots, LysoTracker fluorescence intensity remained unchanged across Lean, Ob4w and FOb mice (**Figure S7 E-H)**.

Together these findings identify impaired lysosomal acidification as a key mechanism driving defective collagenolysis after WL in EpiWAT. This deficiency likely contributes to the accumulation of fibrotic ECM components in the post-obese state, despite suppression of collagen gene expression.

## Discussion

Weight loss, whether achieved through caloric restriction, pharmacological intervention or bariatric surgery, is widely recognized for its ability to improve insulin sensitivity and many metabolic complications associated with obesity. However, whether WAT fully recovers following weight reduction remains incompletely understood. Recent evidence has highlighted the concept of an “obesogenic memory”, whereby prior obesity may leave persistent molecular and cellular alterations despite normalization of body weight and metabolic parameters (23, 44). Our findings extend this concept by demonstrating that initial stage of WL (before weight stabilization) is accompanied by profound remodeling of vVAT and does not restore ECM homeostasis. Thus, our data indicate that the WL phase represents an active remodeling state characterized by progressing structural alterations that extend beyond the simple maintenance/memory of the obese phenotype.

Using the C3H/HeOuj mouse model, a well-established model of obesity-associated vWAT fibrosis characterized by robust ECM deposition (7, 8), we investigated whether obesity-induced ECM alterations are efficiently resolved following WL. This fibrotic phenotype provides the opportunity to examine the mechanisms governing fibrosis and remodeling in the post-obese state, beyond the reversal of adiposity.

Despite improvements in body weight, adiposity and glucose homeostasis, we observed persistent fibrosis in EpiWAT after WL. Importantly, this persistence was not associated with sustained activation of classical fibrogenic programs. Instead, our data indicate that obesity establishes a pro-fibrotic ECM environment that is incompletely resolved after WL. This finding is consistent with our previous observations in human adipose tissue following bariatric surgery (24).

In both experimental models and humans, vWAT fibrosis has been linked to impaired metabolic flexibility and insulin resistance (4). By constraining adipocyte expandability and limiting tissue plasticity (18, 45), fibrosis promotes adipocyte dysfunction (17), suggesting that its incomplete resolution after WL may represent a major obstacle to functional vWAT recovery. The mechanisms maintaining fibrosis after WL appear distinct from those responsible for fibrosis development during obesity. During obesity progression, ECM accumulation is tightly linked to chronic inflammation and the expansion of pro-fibrotic stromal populations, including fibro-inflammatory progenitors (FIPs) (5, 7), and inhibition of fibro-inflammatory pathways effectively reduces vWAT fibrosis in obesity (7, 8). In contrast, post-obese WAT macrophages display reduced expression of genes involved in TNF signaling via the NF-κB pathway, indicating that canonical inflammatory signaling is attenuated after weight loss. Consistent with this, TLR4 inhibition failed to alleviate fibrosis persistence. Although we cannot exclude the contribution of alternative inflammatory pathways capable of maintaining residual NF-κB activity, these findings suggest that persistent fibrosis in the post-obese state is no longer primarily driven by chronic inflammatory signaling.

FIPs represent a subset of PDGFRα-expressing adipose progenitor cells that acquire a pro-inflammatory and pro-fibrotic phenotype during obesity, contributing to ECM production and tissue dysfunction (6, 7). In our study, however, the expression of ECM-related genes within FIP populations was markedly reduced after WL, suggesting that persistent fibrosis is not driven by continued activation of classical collagen-producing pathways. Rather, our findings support a model in which the ECM produced during obesity is poorly resolved after WL. In this context, macrophages and progenitor cells appear to play an important role in ECM turnover. Both populations contributed to collagenolytic activity in vWAT, consistent with evidence from other fibrotic settings, including wound repair, where immune and stromal cells cooperate to regulate ECM deposition and degradation (39, 46). Importantly, despite reduced expression of fibrotic programs after WL, collagen degradation capacity remained impaired in vWAT. To directly assess the contribution of ongoing ECM production, we pharmacologically inhibited ECM synthesis. These interventions did not affect body weight dynamics, although potential influences on food intake or stress-related parameters cannot be fully excluded. While this approach reduced fibrosis during obesity development (7, 8), it failed to prevent fibrosis progression after WL, further highlighting the central role of defective ECM clearance in this context.

Lysosomes are central regulators of ECM remodeling because collagen degradation requires efficient uptake and intracellular processing of matrix components (47). Interestingly, our data revealed an apparent uncoupling between lysosomal gene expression and lysosomal function after WL. While lysosome-associated transcripts were increased in some cell populations, functional assays demonstrated reduced lysosomal activity and impaired collagenolysis. This discrepancy suggests a state of lysosomal stress, where cells attempt to compensate transcriptionally but fail to restore effective degradative capacity (42). Such persistent lysosomal dysfunction may therefore represent a critical bottleneck preventing ECM resolution after WL.

Our findings also suggest that fibrosis persistence after WL is depot-dependent. The impairment in collagen degradation and lysosomal alterations were primarily detected in vWAT, whereas subcutaneous WAT did not exhibit comparable defects. This observation is consistent with the greater susceptibility of vWAT to obesity-induced fibrosis and highlights the importance of considering WAT depot-specific mechanisms when investigating tissue recovery after WL.

Sex-specific differences represent an additional important consideration. Although sexual dimorphism in obesity susceptibility and metabolic responses is well established (29), WAT remodeling in females remain less explored. In our study, female C3H mice developed obesity-associated fibrosis, although to a lower extent than males, but did not display further collagen accumulation after WL. Consistent with this observation, collagen degradation capacity and lysosomal content were preserved in obese females and after WL. These findings suggest that maintenance of ECM turnover mechanisms may limit collagen deposition after WL in female vWAT. However, these observations remain specific to the C3H strain and further studies in additional genetic backgrounds as well as longer weight maintenance periods will be required to determine the generalizability of these mechanisms.

Finally, our findings are consistent with observations in human WAT following bariatric surgery, where progressing ECM deposition have been reported despite improvements in systemic metabolic parameters (24). Although cross-species translation cannot be formally established, the recurrence of similar features across experimental models and human studies suggests that incomplete fibrosis resolution may represent a conserved feature of WAT recovery after WL.

More broadly, our results support the concept that the quality of WAT remodeling after WL may be an important determinant of long-term metabolic health. Although not directly addressed in the present study, our findings raise the possibility that incomplete restoration of WAT architecture could compromise the ability of vWAT to adapt to a subsequent obesogenic challenge. Persistent fibrosis is known to restrict adipocyte expandability and tissue plasticity and may also impair the endocrine functions of vWAT, thereby compromising its ability to maintain systemic metabolic homeostasis. Whether these structural and functional alterations represent a transient phase of post-WL remodeling or persist during long-term weight maintenance remains unknown and warrants further investigation. Taken together, our findings demonstrate that WL reduces obesity-associated inflammatory and fibrotic signaling but does not immediately restore ECM homeostasis in vWAT. Fibrosis persistence after WL likely reflects an imbalance between basal ECM production and its clearance, rather than sustained fibrogenic transcriptional activity alone. These findings highlight the importance of targeting fibrosis resolution, including lysosomal function, rather than solely ECM synthesis, to improve WAT recovery after WL.

## Methods

### Sex as a biological variable

Our study examined male and female animals and sex-dimorphic effects are reported.

### Animals

Mice were housed in groups and maintained on a 12-hour light and dark cycle with ad libitum access to water and standard chow diet (*chow diet*, 3,145kcal/g, 3,1% lipids, *Safe* ref. A04) or high-fat diet (HFD, 5,21kcal/g, 60% lipids, *Research diet* ref. *D12492)*. C3H/HeOuJ (C3H) mice were obtained from Charles River laboratories (JAX:000635). For TAK242 infusion (**Figure S5**), mice were fed HFD for 4 weeks and then implanted intraperitoneally with osmotic minipumps (model 1004, Alzet) filled either with a vehicle solution of DMSO/ ethanol/H_2_O (5/1.5/3.5) or TAK242 (6587 Tocris) to continuously infuse 350 μg/kg/day for 4 weeks, as previously published (8). For treatment with nintedanib and celecoxib, C3H males were treated with vehicle (Carboxymethyl cellulose 0.25%), or nintedanib (100mg/kg) combined with celecoxib (20mg/kg) by daily oral gavage, five times a week (7). Body composition was measured by nuclear magnetic resonance (NMR) on conscious animals using a Bruker analyzer (Minispec LF90). All animal procedures were reviewed and approved by local and national comities, and conducted in accordance with the Guide for the Care and Use of Laboratory Animals published by the European Commission Directive 86/609/EEC.

### Glucose metabolism assessment

For assessment of glucose tolerance test, mice were fasted for 4 hours prior to intraperitoneal glucose injection at a dose of 2 grams per kg of body weight. Glycemia was measured with a glucometer (Accu-Check, Roche) at baseline and 15, 30, 45, 60, 90 minutes. Glucose levels were monitored up to 90 min, as differences between experimental groups were already evident within this time frame and this duration is sufficient to capture glucose excursion and assess glucose tolerance (48). For insulin tolerance test, fasted mice received 1U/kg of insulin and glycemia was measured at baseline and 15, 30, 45, 60 and 90 minutes.

### Collagen content

Hydroxyproline content was measured using a hydroxyproline colorimetric assay (Sigma). Briefly, frozen fat is weighted, distributed in sealed tubes containing 6 N HCl (10 μL of HCl/mg of WAT) and heated overnight at 110°C. For AT, the entire frozen fat pad is homogenized to obtain a homogenate powder before weighing the required amount for analysis. Five microliters are evaporated before incubation with chloramine-T and p-di-methyl-amino-benzaldehyde (DMAB) at 60°C for 90 min. The absorbance is read at 560 nm and the concentration is determined using the standard curve created with purified hydroxyproline.

### mRNA extraction, reverse transcription and quantitative PCR

RNAs from adipose tissue samples were prepared using the RNeasy Tissue kit (Qiagen) and cDNAs were synthesized with M-MLV reverse transcriptase (Promega). Quantitative PCR analyses were performed using the StepOne system (Applied Biosystems) with Fast SYBR Master Mix (Life Technologies). For each sample, *18s* and *Rplp0* were used as housekeeping genes. Relative fold changes in gene expression were calculated by the ΔΔCt method using the equation 2^(-ΔΔCt)^. All primer sequences are available upon request.

### mRNA sequencing, data preprocessing and analysis

Total RNA preparation was performed from 20000 sorted cells (adipose progenitors or macrophages) using the RNeasy Micro Kit (Qiagen). cDNA libraries were generated using the KAPA mRNA HyperPrep kit (Roche). RNA-Seq libraries were sequenced on an Illumina NovaSeq 6000 (30 million reads per sample). RNAseq analysis was completed using the Eoulsan 2.5 pipeline. The minimap2 version 2.18 index was used to map raw reads to the genome and data normalization was performed with DESeq2. A linear model for RNAseq analysis (LIMMA) was conducted to select differentially expressed genes with a 1.5-fold change cutoff between at least two conditions. Annotated genes with a count mean over 10 in at least one condition and a coefficient of variation of more than 0.5 between at least two conditions were retained. Analysis of the enriched pathways were conducted with GSEA (http://software.broadinstitute.org/gsea/index.jsp) (49) and GenePattern platforms (http://www.broadinstitute.org/cancer/software/genepattern/) (50), respectively. RNA sequencing data have been deposited with the data set identifier E-MTAB-15063.

### Immunofluorescence on paraffin-embedded AT sections

Mouse epididymal AT sections were cut to a thickness of 5 µm using a microtome and then incubated with different antibodies: Anti-perilipin 1 antibody (GP33 Progen; 1 :100 dilution) ; IHC: Nε-(Υ-L-Glutamyl)-L-Lysine Isopeptide Antibody (81D4) (Covalab 00012721 monoclonal antibody; 1 :100 dilution) ; Cross Linked C-Telopeptide Of Type I Collagen (CTXI) (Cloud-clone corp MAA665Hu2 monoclonal antibody; 1 :200 dilution). The primary antibodies were detected using the Alexa fluor 647 Tyramide SuperBoost amplification kit (Invitrogen, B40936) according to the manufacturer’s instructions.

### Sirius red staining for fibrosis

Sections of epididymal and inguinal adipose tissue from mice, embedded in paraffin, were cut at a thickness of 5 µm using a microtome. After the deparaffinization steps in alcohol solutions, slides were incubated for 1 hour in picrosirius red. Slides were mounted with Eukitt mounting medium (Sigma-Aldrich 03989). Images were captured using a Eclipse TiE Nikon Inverted microscope with a polarizer (51–53). Images were analyzed with ImageJ (National Institutes of Health by Wayne Rasband).

### Ex vivo Lipolysis

WAT explants were incubated in Dulbecco’s Modified Eagle’s Medium (DMEM, ThermoFisher, ref: 21063029) with or without 10^-8^M CL 316-243 (Sigma Aldrich, 138908-40-4) for 90 minutes at 37°C as 100 mg of tissue/mL. Glycerol concentration released into the medium was determined with a colorimetric assay kit (Sigma Aldrich, MAK117) following the manufacturer’s instructions. Glycerol was normalized per cell number (54).

### Flow cytometry

Mouse antibodies were purchased from Biolegend : CD45 (30-F11), Ly-6C (HK1.4), CD31 (390), CD140a (APA5), CD9 (MZ3), CD64 (X54-5/7.1); from Thermo Fisher Scientific : PDPN (8.1.1); from BD Biosciences Ki-67 (B56). Cell suspensions were stained with appropriate antibodies for 30 min on ice. Draq7 dye was used to exclude dead cells. Intracellular Ki-67 staining was performed using the Foxp3 staining kit from Thermo Fisher Scientific. Data were acquired on a BD LSRFortessa™ flow cytometer (BD Biosciences) and analyzed with FlowJo software (Tree Star). To calculate absolute counts, a fixed number of beads was added to each tube as previously described (55). Cell sorting was performed on a BD FACSAria II™ cell sorter.

### In situ zymography

WAT explants were incubated in DMEM containing 10% serum with or without 20 μg/mL of DQ collagen I (ThermoFisher Scientific, D12060) at 37°C for 1 hour. After incubation, tissues were washed with cold PBS and then digested in 1 mg/ml of collagenase A (Roche, ref: 110887933001) at 37°C for 30 minutes. Cells from the stromal vascular fraction (SVF) were analyzed by flow cytometry.

### Lysosome quantification

SVF was incubated with LysoTracker (ThermoFisher) for 30 minutes at 37°C and washed in cold PBS. Cells were stained with antibody as above and were then analyzed by flow cytometry. Cells isolated from the adipose tissue stromal vascular fraction (SVF) exhibit variable levels of autofluorescence signal. To account for this nonspecific background, LysoTracker analyses were performed using matched unstained controls for each individual sample (i.e., with and without LysoTracker staining), allowing the LysoTracker-positive population to be defined while accounting for sample-specific autofluorescence.

### Preparation of tissue for LC-MS

The tissue was suspended in an extraction buffer (ABC 50 mM/DTT 10mM/SDC 2%) and heated for 5 min at 95°C to reduce and extract proteins. IAM (50mM final) was added to alkylate the free cysteine residues at r.t. for 20 min. The protein solution was diluted in ABC 50 mM to reach SDC 0.5% final concentration before adding the mix trypsin-LysC and incubated overnight at 37°C with constant shaking. The digests were centrifuged to recover the supernatant. A volume of TFA 10% was added to quench the trypsin and to precipitate SDC as previously described (56). The peptides were desalted on Zip-Tip C18 (Millipore/Merck). After removal of acetonitrile with a SpeedVac (Savant ISS110), the desalted peptides were resuspended in solvent A’ (H2O/ACN 98:2 (v/v), 0.1% TFA) first in 15µl. Nano LC-MS/MS analyses 2µl of each sample was injected on a reversed-phase trap column (C18 Acclaim PepMap100, 5 μm, 300 μm i.d. × 5 mm) and eluted to a reversed-phase column (C18 Waters, 2 μm, 75 μm i.d. × 25 cm) at a constant flow rate of 300 nL/min, with a gradient of 2–44% of solvent B (H2O/ACN 10:90 (v/v), 0.1% FA) in solvent A (H2O/ACN 98:2 (v/v), 0.1% FA) over 123 min. The LC was coupled to a Orbitrap Eclipse mass spectrometer (Thermo Fisher Scientific) equipped with a nano-electrospray ion source. Top S DDA MS experiments consisted of one MS survey scan in the Orbitrap cell (resolution of 60000, scan range 375–1500 m/z, AGC target 3e6, maximum injection fill time 60 ms) followed by the MS/MS scans of the most intense precursors in HCD mode (dynamic exclusion 30 s, no singly charged precursors, resolution 15000, NCE 28, AGC target 1e4, maximum injection time 60 ms, isolation window 2 m/z).

Validation and processing. Data were processed using Andromeda in Maxquant 2.0.3.0 against the Mouse SwissProt database/ human Swissprot Database (released May 2025) with the following parameters:10 ppm MS mass tolerance, 0.6 Da MS/MS mass tolerance, enzyme trypsin, carbamidomethylation of cysteine (C), acetylation (Protein N-term) and oxidation of methionine (M) as variables modifications, maximum 2 miscleavages and 1% false discovery rate (FDR). Protein annotations were given by the Uniprot database and the matrisome database build by the Hyne Lab (http://matrisomeproject.mit.edu/)(57).

## Data availability

Data are available in public repositories. RNA sequencing data have been deposited with the data set identifier E-MTAB-15063. Proteomic data have been deposited on PRIDE with the data set identifier PXD064522. Supporting data values associated with the main manuscript and supplement material are provided in the “Supporting data values” file.

## Statistical analysis

Data are expressed as mean ± SEM. Normality of continuous variables was assessed using the Shapiro-Wilk test. If the data did not follow a Gaussian distribution, a natural logarithm transformation or a square root transformation (for data with values equal to zero) was applied. Outliers were identified using the ROUT method (false discovery rate < 1%). Comparisons between two groups were performed using 2-tailed Student’s t-test. Where indicated, one-way ANOVA followed by multiple comparison tests was performed. Statistical analyses were conducted using R and GraphPad Prism. A P value less than 0.05 was considered significant and represented as follows *p<0.05 ; **p<0.01 ; ***p<0.001 (unless otherwise indicated).

## Study approval

All animal studies have been approved by the relevant institutional review boards (Comité Charles Darwin – CE2A005, Paris, France) and the French ministry of higher education and research (Ministère de l’enseignement supérieur et de la recherche) under the reference number APAFIS#54012

## Author contributions

**Conceptualization,** E. L. G., G. M., K. C. **Methodology,** Ad. F., E. L. G., F. A., G. M, I. D., J. V., K. C., L. G, M. O, S. L., S. V. **Investigation** Ag. F., A. O., A. B., A. S., C. B., Ci. R., Cl. R, Ch. R., F. M., I. H., J.B.A., L. B., L. T., S. D. **Original draft**, E.L.G., G.M., K.C. All authors reviewed, read and approved the manuscript.

## Acknowledgments

We would like to express our gratitude to all the nursing and medical staff. We would like to thank all of the personnel of the animal facility (UMS28) and all the personnel of the flow cytometry platform (CyPS) of Sorbonne University (Paris-France). We thank ICM platforms (ICM Quant, iGenSeq and Histomics) as well as ‘Service d’anatomie et cytologie pathologiques’ at Pitié-Salpêtrière Hospital for their help with histological sample preparation and analysis.

