## Supplemental figures for "A Weight Loss Fibrotic Response in Visceral Fat Is Driven by Defective Lysosome Dependent Collagenolysis"

### Supplemental figures and legends

**Figure S1. Normalized glucose metabolism and IngWAT features in FOb mice.**

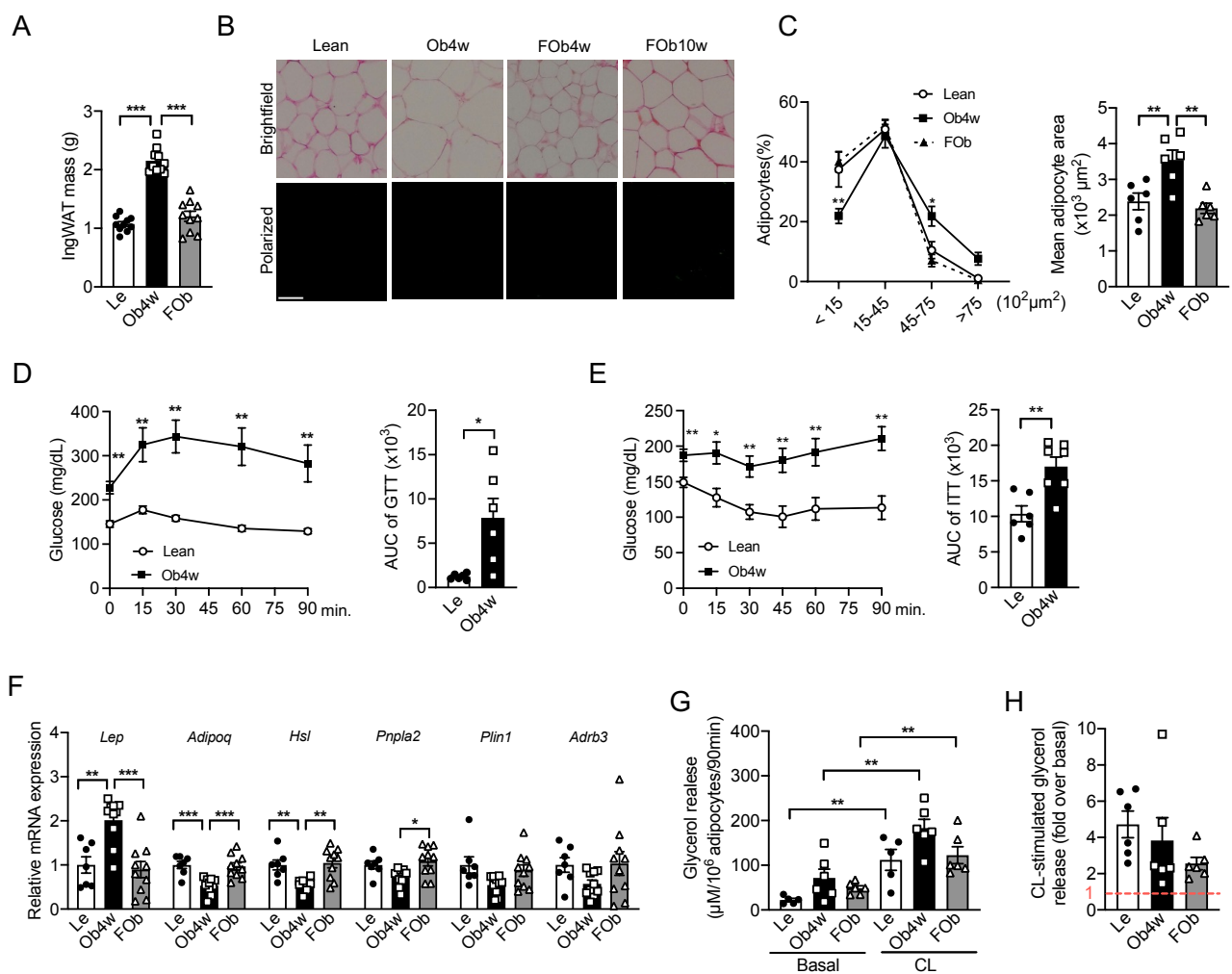

(A) IngWAT mass in lean (Le), obese (Ob4w) and formerly obese (FOb) mice (n = 10). (B) Representative picosirius red-stained paraffin sections of IngWAT from Le, Ob4w and FOb C3H mice, imaged under brightfield and polarized light (scale bar, 100 μm). We analyzed two FOb groups, both fed a high-fat diet (HFD) for 4 weeks, then switched to a chow diet for either 4 weeks (named FOb or FOb 4w) or 10 weeks (FOb 10w). (C) Adipocyte area distribution and mean adipocyte size in Le, Ob4w and FOb mice (n = 5–6). (D) Glucose tolerance test (GTT) in age-matched Le and Ob4w mice at week 4 (n = 6). The area under the GTT curves (AUC) is shown. (E) Insulin tolerance test (ITT) in age-matched Le and Ob4w mice at week 4 (n = 6). The area under the ITT curves (AUC) is shown. (F) Relative mRNA expression of adipocyte markers in IngWAT measured by quantitative PCR (n = 7–10). (G) Basal and CL-316,243 (CL)-stimulated glycerol release from IngWAT explants, normalized to adipocyte number, compared between Le, Ob4w and FOb mice (n = 5–6). (H) Fold change in CL-induced lipolysis over basal glycerol release in IngWAT explants from Le, Ob4w and FOb mice (n = 6). (A, C, F–H) Statistical significance was assessed using one-way ANOVA followed by Tukey's multiple comparisons test. (D, E) Student's t-test.

**Figure S2. Glucose tolerance and visceral white adipose tissue hydroxyproline content in female mice before and after weight loss**

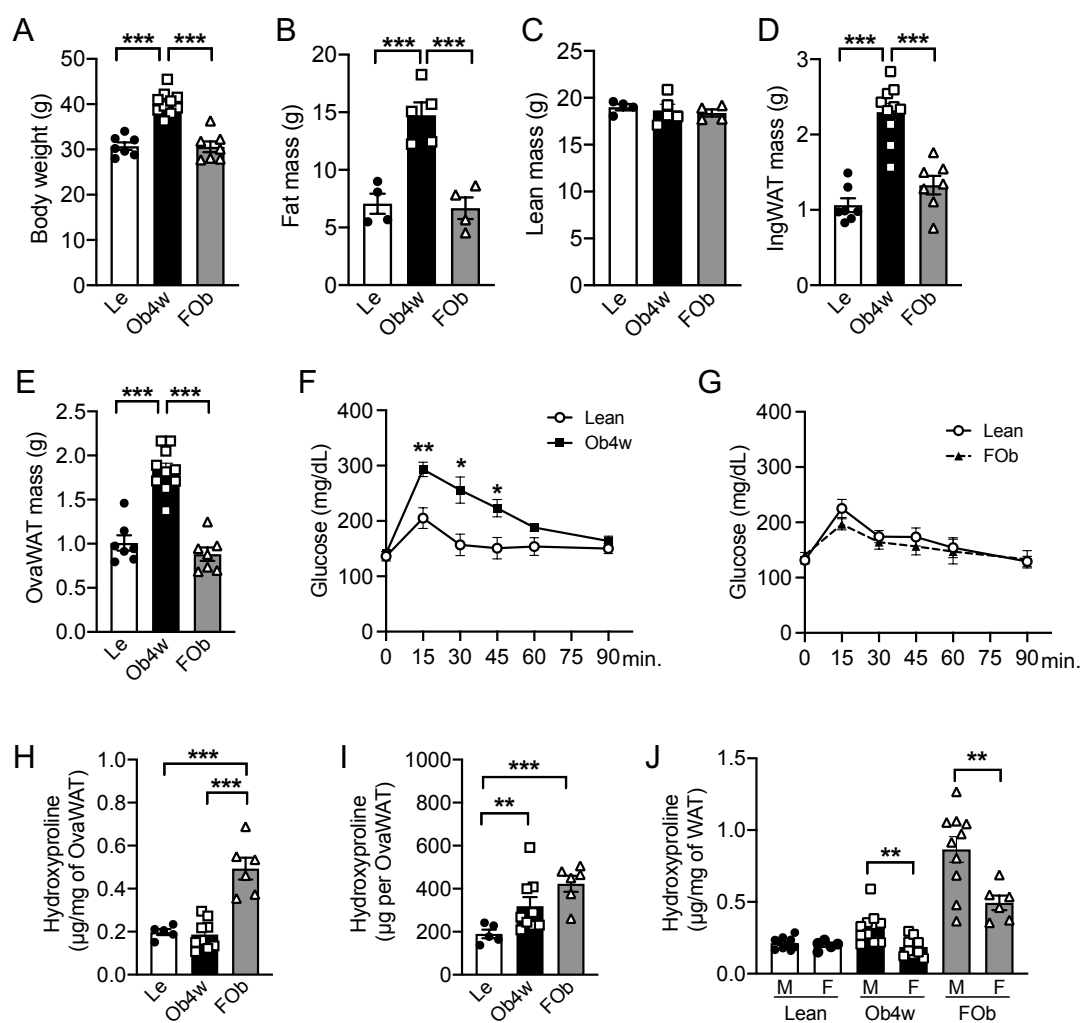

(A) Body weight (n=7-10), (B) fat mass (n=4-5), (C) lean mass (n=4-5), (D) IngWAT and (E) OvaWAT masses (n=7-10) in Le, Ob4w and FOb females (F) Glucose tolerance test (GTT) in age-matched Le and Ob4w mice after 4 weeks of dietary intervention (n = 4). (G) GTT in age-matched Le and FOb mice after 8 weeks of dietary intervention (n = 5-6). (H) Hydroxyproline content in OvaWAT normalized to fat pad mass (μg/mg) and (I) total hydroxyproline content in total OvaWAT pad (n=5-9). (J) Hydroxyproline content in EpiWAT (for male, M) and OvaWAT (for female, F) normalized to fat pad mass (μg/mg) in lean, Ob4w and FOb (n=5-10). (A-E, H, I) Statistical significance was assessed using one-way ANOVA followed by Tukey's multiple comparisons test. (F, G, J) Student's t-test.

90 **Figure S3. Matrisome proteins in EpiWAT from Lean (Le), Obese (Ob8w) and Formerly obese**
91 **mice (FOb).**

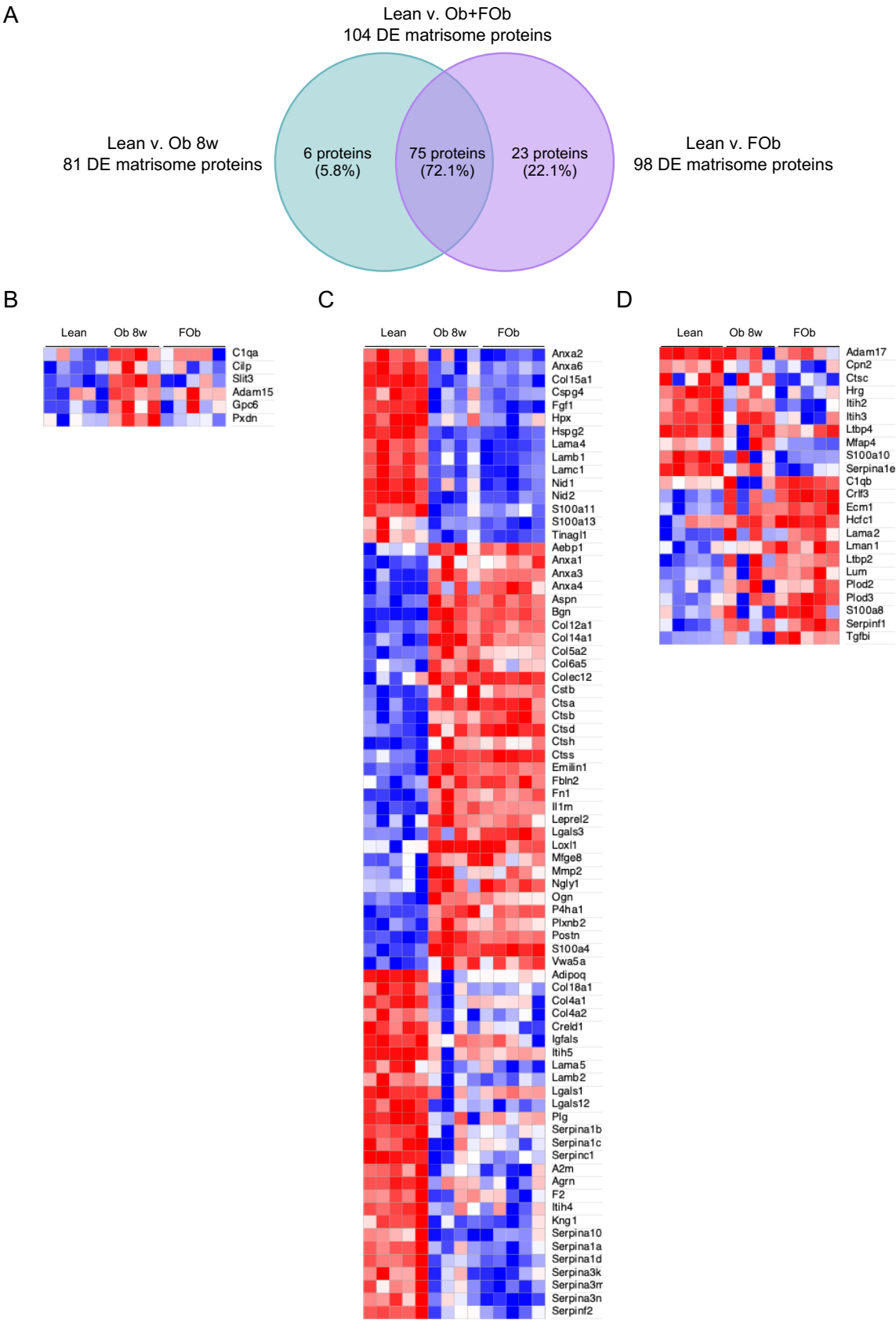

92 (A–D) Venn diagrams and expression heatmaps showing differentially expressed (DE) matrisome
93 proteins (n=4-5). (A, C) Overlapping DE proteins between Lean vs. Ob8w and Lean vs. FOb
94 comparisons. (A, B) DE proteins specific to the Lean vs. Ob8w comparison. (A, D) DE proteins
95 specific to the Lean vs. FOb comparison. (A–D) A Limma analysis was conducted to select
96 differentially expressed proteins between Le vs. FOb and Le vs. Ob8w

**Figure S4. Gating strategy to define macrophages and progenitor cells in adipose tissue**

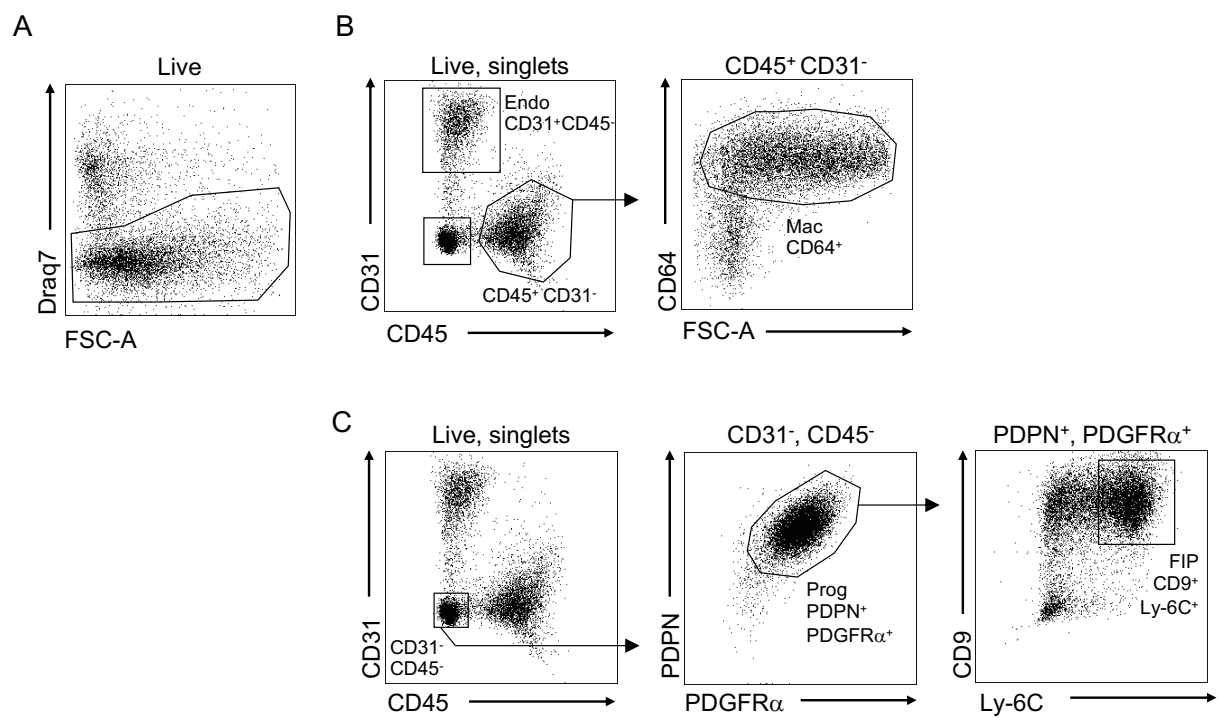

**(A)** Cells were first gated on live cells using a viability dye to exclude dead cells. **(B)** Endothelial cells (Endo) were identified as CD31<sup>+</sup> CD45<sup>-</sup> cells. For macrophages, CD45<sup>+</sup> CD31<sup>-</sup> immune cells is then followed by selection of macrophages (Mac) identified as CD64<sup>+</sup> cells. **(C)** Gating strategy used for the selection of total progenitors (Prog, PDGFRα<sup>+</sup> PDPN<sup>+</sup> CD31<sup>-</sup> CD45<sup>-</sup>) and CD9<sup>+</sup> Ly-6C<sup>+</sup> fibro-inflammatory progenitors (FIP).

**Figure S5. Weight loss associated collagen deposition is not abrogated with TAK 242 or with** **Nintedanib+Celecoxib.**

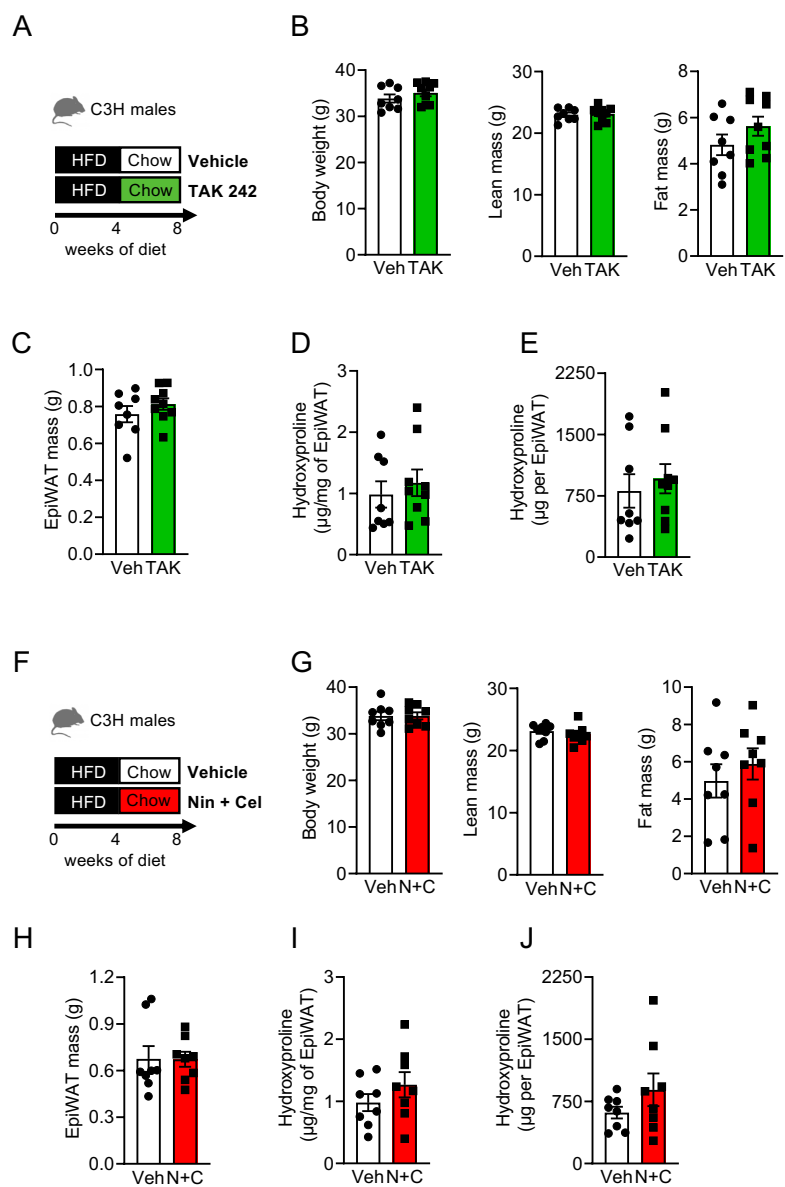

(A) Experimental design showing dietary interventions (chow or high-fat diet, as indicated) and the timeline for obesity induction and weight loss. Mice were implanted intraperitoneally with osmotic pumps containing either vehicle (white) or TAK242 (green), which were infused continuously for four weeks during the WL period. (B) Body weight, body composition (lean and fat mass) and (C) EpiWAT mass. (D–E) Fibrosis in EpiWAT from formerly obese (FOb) mice treated with vehicle or TAK242 was quantified by measuring (D) Hydroxyproline content normalized to fat pad mass (μg/mg) and (E) Total hydroxyproline content per fat pad (n = 8–9). (F) Experimental design showing dietary interventions (chow or high-fat diet, as indicated) and the timeline for obesity induction and weight loss. Mice were treated with Veh or with the combination Nintedanib and Celecoxib (Nin+Cel) during the WL period. (G) Body weight, body composition (lean and fat mass) and (H) EpiWAT mass. (I–J) Fibrosis in EpiWAT from formerly obese (FOb) mice treated with vehicle or Nin+Cel was quantified by measuring (I) Hydroxyproline content normalized to fat pad mass (μg/mg) and (J) Total hydroxyproline content per fat pad (n = 8). (B–E, G–J) Statistical significance was assessed using Student's t-test.

Expression heatmap showing genes from selected pathways (**Figure 3G, H**) that are enriched in EpiWAT macrophages (Macs) from either obese (Ob4w) or formerly obese (FOb) mice. Data from lean (Le), Ob4w and FOb mice are presented.

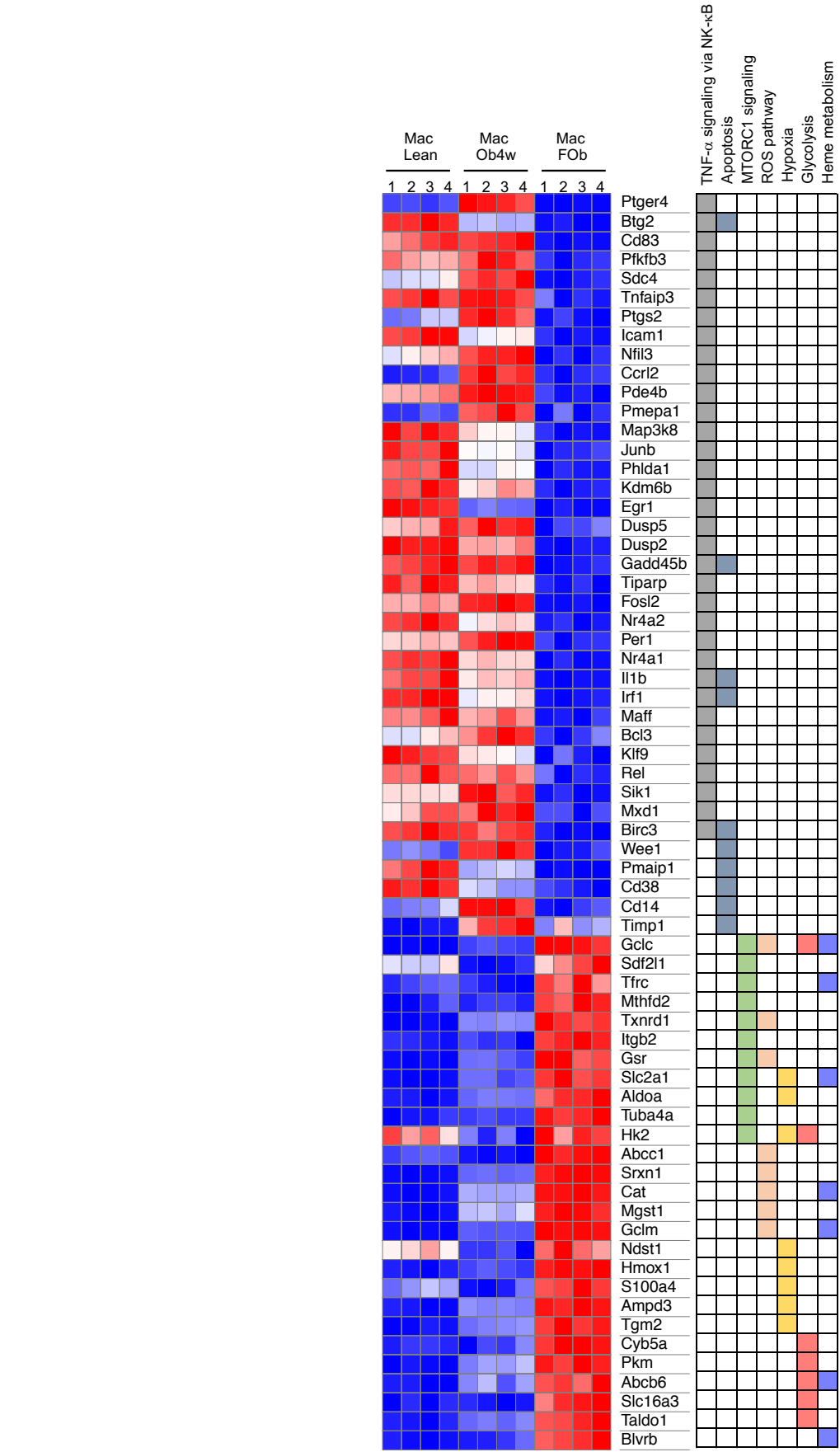

**Figure S7. DQ-Collagen I degradation and lysotracker staining in macrophages and progenitors from IngWAT of males and OvaWAT of females**

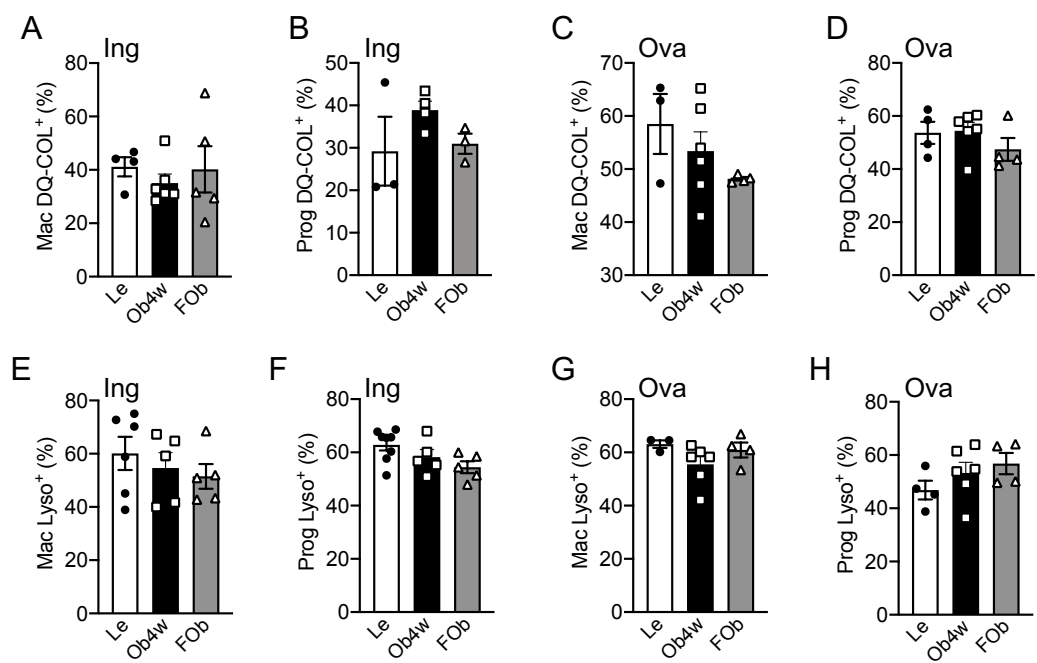

(A, B) Percentage of DQ-COL1<sup>+</sup> cells in (A) Macs (n=4-6) and (B) Progs (n=3-4) isolated from IngWAT of lean (Le), obese (Ob4w) and formerly obese (FOb) C3H males. (C, D) Percentage of DQ-COL1<sup>+</sup> cells in (C) Macs (N=3-6) and (D) Progs (n=4-6) isolated from OvaWAT of lean (Le), obese (Ob4w) and formerly obese (FOb) C3H females. (E, F) Percentage of LysoT<sup>+</sup> in (E) Macs (n=5-6) and (F) Progs (n=5-8) isolated from IngWAT SVF of Le, Ob4w and FOb males. (G) Percentage of LysoT<sup>+</sup> in (G) Macs (n=3-6) and (H) Progs (n=4-6) isolated from OvaWAT SVF of Le, Ob4w and FOb females. (A-H) Statistical significance was assessed using one-way ANOVA followed by Tukey's multiple comparisons test.

**Figure S8. Relative mRNA expression of ECM remodeling enzymes and genes involved in** **collagen uptake.**

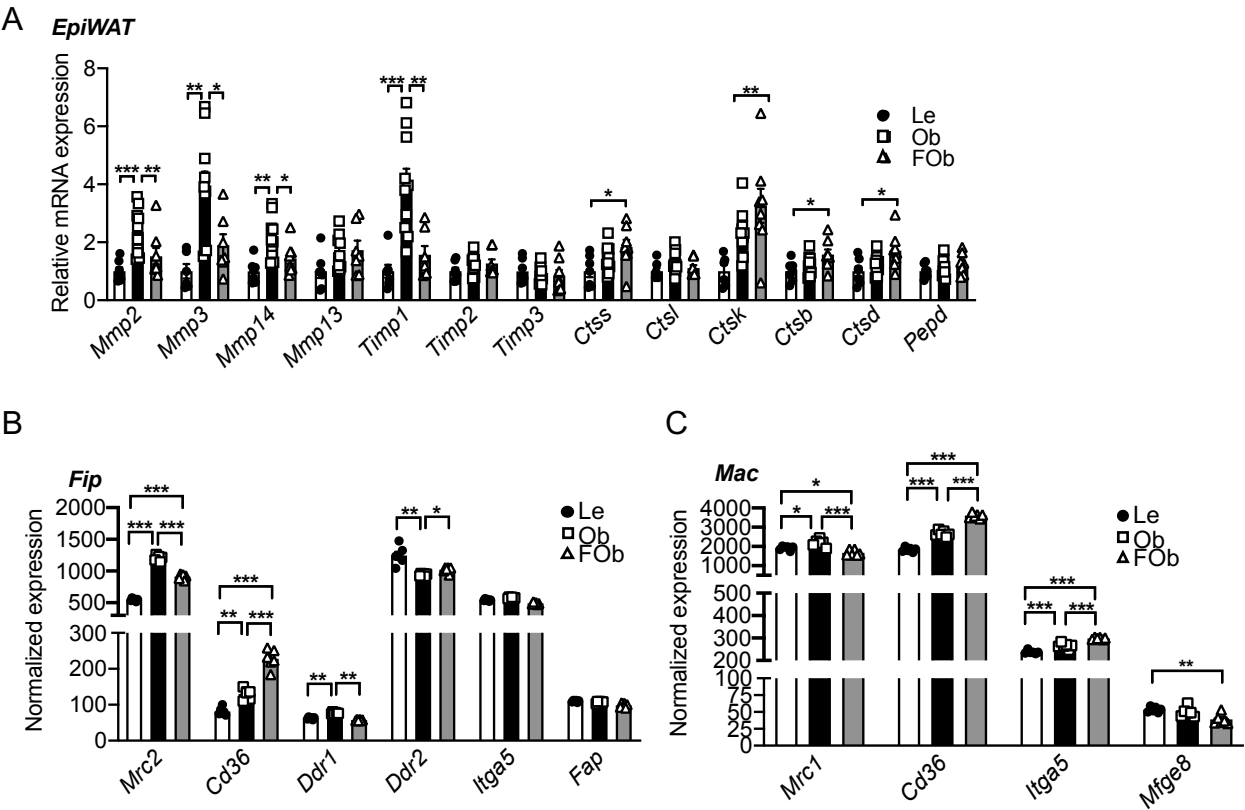

(A) mRNA expression levels of metalloproteinases (*Mmp2*, *Mmp3*, *Mmp14*, *Mmp13*), tissue inhibitors of metalloproteinases (*Timp1*, *Timp2*, *Timp3*), cathepsins (*Ctss*, *Ctsl*, *Ctsb*, *Ctsd*), and peptidase D (*Pepd*) in EpiWAT from lean (Le), obese (Ob4w), and formerly obese (FOb) C3H mice, measured by quantitative PCR (n=6–10).
(B, C) mRNA expression of genes involved in collagen uptake analyzed in (B) FIPs (n=6) and (C) Macs (n=4), based on RNA sequencing of FACS-sorted cells from Le, Ob4w, and FOb EpiWAT of C3H mice. (A–C) Statistical testing was performed using one-way ANOVA followed by Tukey’s multiple comparisons test.

**Figure S9. The lysosome pathway is upregulated in macrophages (Macs) and FIPs from obese** **(Ob) and formerly obese (FOb) mice compared to lean mice.**

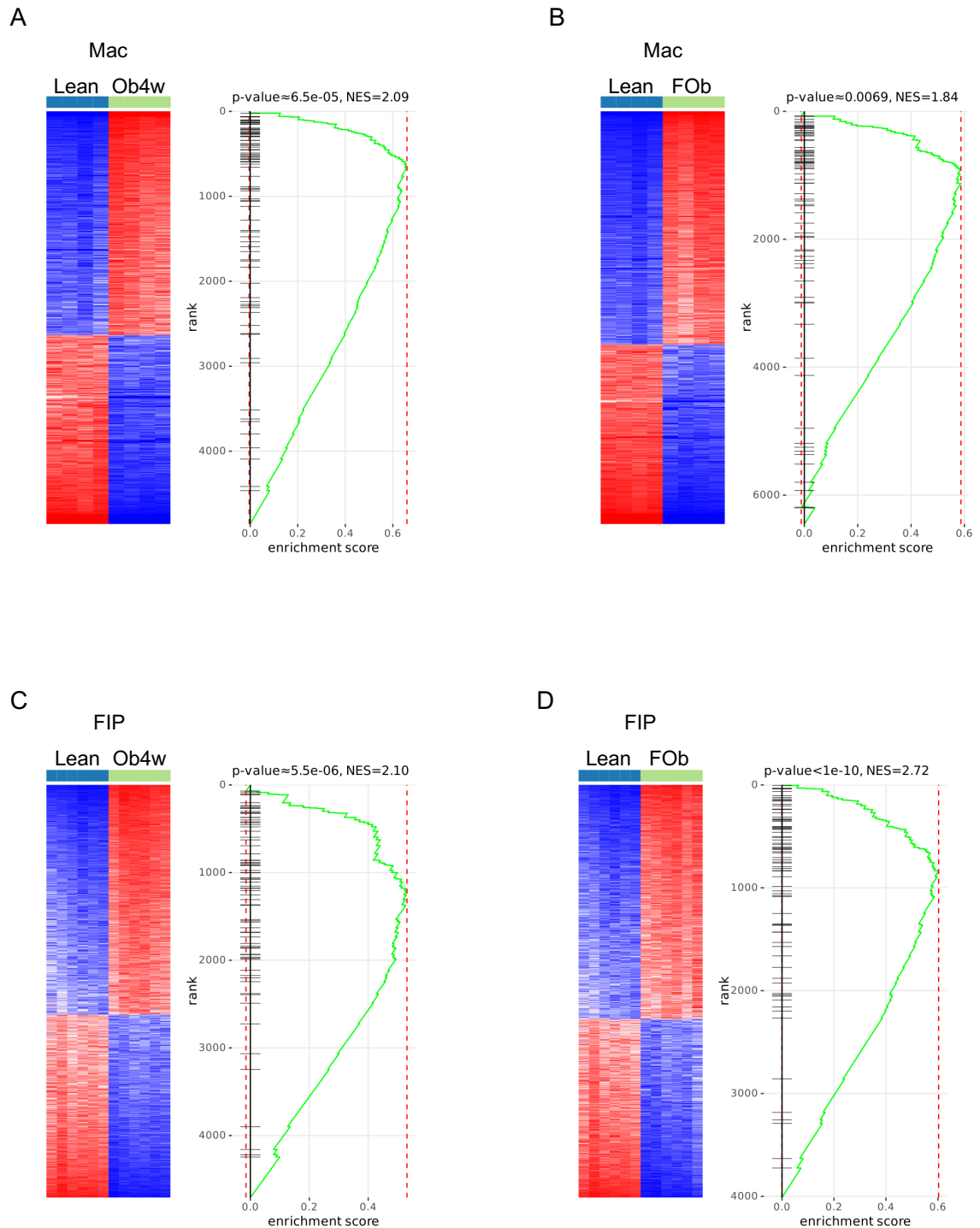

Gene set enrichment analysis (GSEA) of genes from the lysosome KEGG pathway in Macs and FIPs from (A, C) lean vs. obese, and (B, D) lean vs. formerly obese (FOb) mice. Differentially expressed genes were identified using LIMMA analysis, and GSEA was performed using the Phantasus platform on RNA-sequencing data from Macs and FIPs isolated from EpiWAT of lean, obese, and FOb C3H mice.
